# Influence of activation pattern estimates and statistical significance tests in fMRI decoding analysis

**DOI:** 10.1101/344549

**Authors:** Juan E. Arco, Carlos González-García, Paloma Díaz-Gutiérrez, Javier Ramírez, María Ruz

## Abstract

The use of Multi-Voxel Pattern Analysis (MVPA) has increased considerably in recent functional magnetic resonance imaging studies. A crucial step consists in the choice of methods for the estimation of responses and their statistical significance. However, a systematic comparison of these and their adequacy to predominant experimental design is missing.

In the current study, we compared three pattern estimation methods: Least-Squares Unitary (LSU), based on run-wise estimation, Least-Squares All (LSA) and Least-Squares Separate (LSS), which rely on trial-wise estimation. We compared the efficiency of these methods in an experiment where sustained activity had to be isolated from zero-duration events as well as in a block-design approach and in an event-related design. We evaluated the sensitivity of the *t*-test in comparison with two non-parametric methods based on permutation testing: one proposed in Stelzer et al. (2013), equivalent to performing a permutation in each voxel separately and the Threshold-Free Cluster Enhancement (Smith and Nichols, 2009).

LSS resulted the most accurate approach to address the large overlap of signal among close events in the event-related designs. We found a larger sensitivity of Stelzer’s method in all settings, especially in the event-related designs, where voxels close to surpass the statistical threshold with the other approaches were now marked as informative regions.

Our results provide evidence that LSS is the most accurate approach for unmixing events with different duration and large overlap of signal, consistent with previous studies showing better handling of collinearity in LSS. Moreover, Stelzer’s potentiates this better estimation with its larger sensitivity.

## 1. Introduction

Multi-Voxel Pattern Analysis (MVPA) has become a widely used technique in functional Magnetic Resonance Imaging (fMRI) studies. MVPA employs brain activation patterns to discriminate between experimental conditions of interest. This can be considered as a classification problem, where the classifier uses the features contained in the patterns (e.g. the voxels in the image) to learn the relationship between them and the experimental conditions. Then, based on this learning, the classifier predicts the experimental conditions to which new images belong using only their activation patterns. Since the classifier uses this information as input, the result of the classification process depends on the quality of the patterns and the way they are estimated. The sluggishness of blood-oxygen-level-dependent (BOLD) signal adds difficulty to this classification endeavor: during an experimental condition, the BOLD signal increases about 2 seconds after neural activity, peaking at about 5-8 seconds later and returning to baseline approximately at 20 seconds (Logothetis and Wandell, 2004). In block designs, where an experimental condition is presented continuously for an extended time interval, isolating the relevant signal is relatively straightforward. This is similar to slow-event related designs where the inter-stimulus-interval (ISI) is longer than the duration of the BOLD. However, when the ISI is short (such as in rapid event-related designs), there is a large overlap between trials, which complicates the estimation of the contribution of each one of them to the combination of individual hemodynamic responses (see Figure 1).

**Figure 1:**
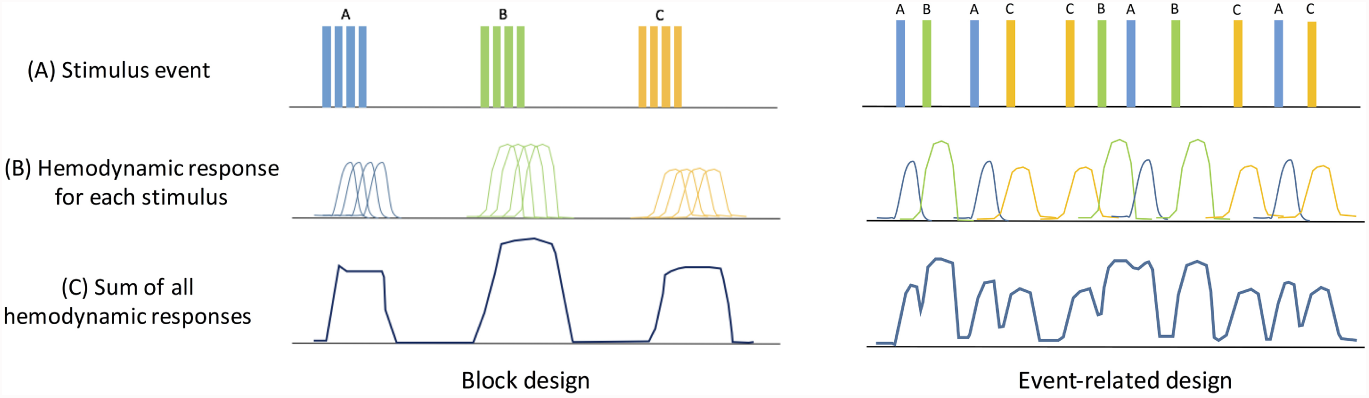
Schematics of two different fMRI designs: block and event-related. The first row corresponds to the timing of event onsets. In block designs, several stimuli of the same condition are presented consecutively, in what is known as epoch or block, and different conditions usually alternate in time, so relatively large signal changes are measured. In event-related designs, interleaved short-duration stimuli are employed. Given the delayed nature of the BOLD signal, the data produced by different stimuli overlaps, and thus extracting the signal caused by each one of them becomes more difficult.

Most fMRI analyses use linear convolution models like the General Linear Model (GLM) to extract estimates of responses to different event types (Friston et al., 1998), where the model estimation is carried out voxelwise and the BOLD time series is the dependent variable. The parameters of the GLM are computed by minimizing the squared errors across scans between the timeseries that is predicted, guided by information of the fMRI experiment like stimulus onsets and assuming the shape of the BOLD response and the noise in the data.

Equation (1) shows mathematically how this estimation is performed:
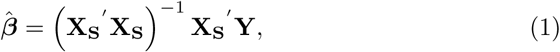

where **Y** is the vector of the BOLD response time series, **X_S_** is the design matrix and *β* is the vector of activation estimates.

Previous studies have explored different methods to obtain activation estimates (also known as beta weights or beta maps) in event-related designs (Abdulrahman and Henson, 2016; Mumford et al., 2012). The most common is the so-called ‘Least-Squares Unitary’ (LSU), in which all trials of the same type (e.g. experimental conditions) are collapsed into one single regressor, and trial variability is relegated to the GLM error term. Other studies have focused on obtaining single-trial parameter estimates. The most straightforward approach is known as beta-series regression (Rissman et al., 2004), in which a different regressor is used for each trial. Following the notation in Mumford et al. (2012), we from now on denote it as ‘Least-Squares All’ (LSA). Figure 2 shows a visual representation of how these two methods work. For two different stimuli (e.g. a letter and a face), LSU estimates the contribution to the hemodynamic signal of each condition along the run, whereas LSA estimates it trial-by-trial. When trials have short ISI, the regressors become highly correlated, which can inflate the variance of the resulting parameter estimates and the subsequent classification accuracies (Mumford et al., 2014). To address this drawback, Turner (2010) introduced an alternative method known as ‘Least-Squares Separate’ (LSS), based on iteratively fitting a new GLM for each trial. There are different variants on this approach depending on the number of regressors defined. In the simplest one, LSS-1, there is a parameter for the target trial and another single nuisance parameter for the rest (see Figure 3). In LSS-2, each GLM includes three regressors: the first one, for the target trial; the second for the rest of the trials of the same type as the target, whereas the third is used for the trials of a different type. It is thus possible to define as many nuisance parameters as trial types (e.g. LSS-N), although LSS-1 (from now on, LSS) is commonly used due to its simplicity and high performance (Abdulrahman and Henson, 2016; Turner et al., 2012).

**Figure 2:**
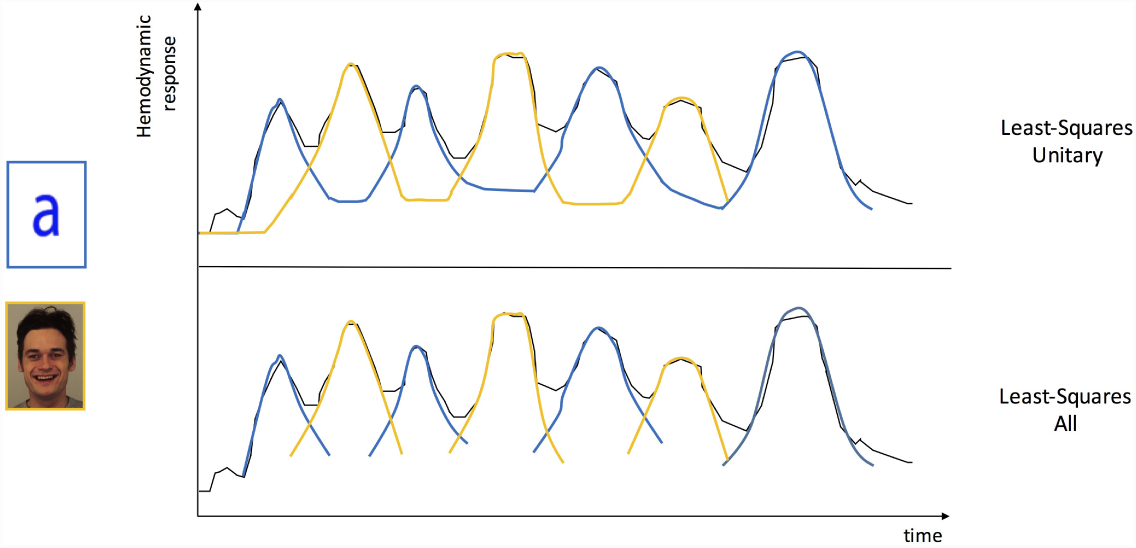
Comparison of two different approaches for pattern estimate. At the top of the figure, LSU, in which all the trials of the same type for each run are collapsed into the same regressor. At the bottom, LSA, based on estimating one model in which each event is modelled as a separate regressor. LSU can yield less noisy estimates because of the averaging of all the stimuli of the same type within a run, but the amount of resulting estimates to train the classifier with is limited to the number of runs the experiment is divided into.

**Figure 3:**
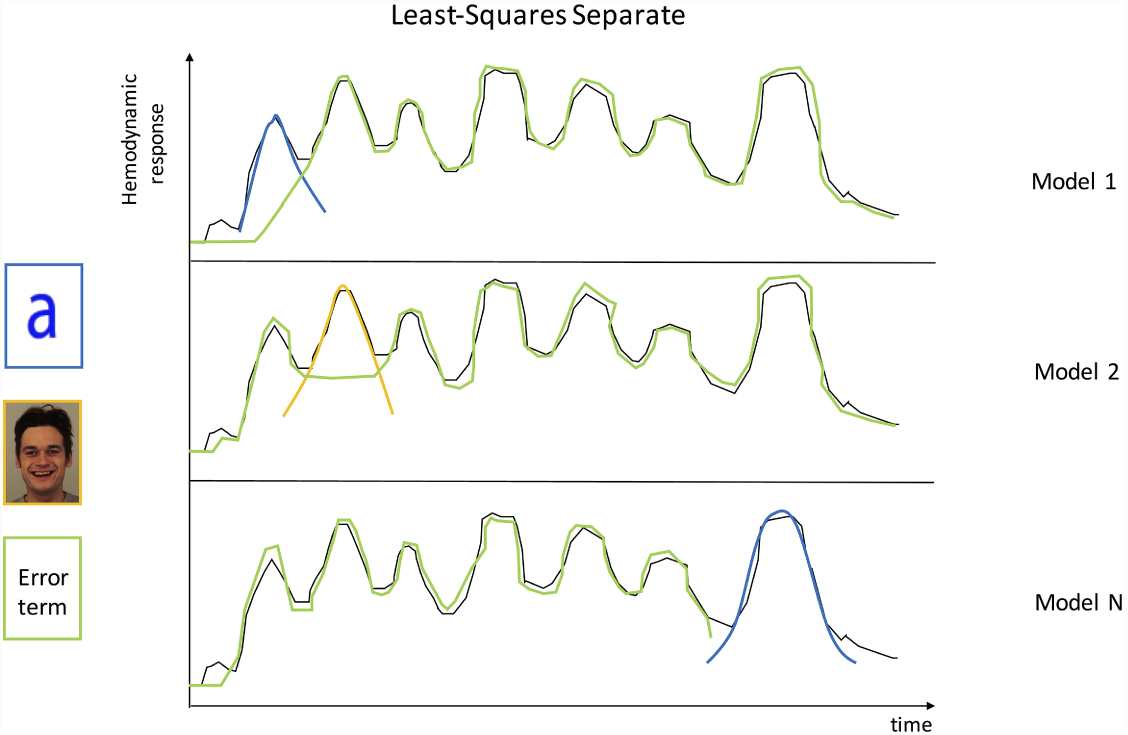
LSS iteratively fits a new GLM for each unique event with two predicted BOLD time courses: one for the target event and a nuisance parameter estimate which represents the activation for the rest of the events. LSS estimates as many models as the total number of regressors, and in each one only two of them are included: one for the event of interest and a nuisance parameter estimate which stands for the activation for the rest of the events.

The advantages of single-trial estimates are reflected in the fields of neuroscience and also machine learning. Regarding the first one, a good example is the study of working memory, where classic models assume that information is stored via persistent neural activity (Sreenivasan et al., 2014). Whereas averaging across trials may cancel out the noise and improve the signal-to-noise ratio, trial-wise averaging may also remove important coding signals (e.g. Stokes and Spaal, 2016). From the machine learning standpoint, estimating one beta map per trial yields a larger number of images to train the classifier, which improves generalization. Pereira et al. (2008) discussed the important tradeoff between having many noisy examples (e.g. one per trial) or fewer, cleaner ones (e.g. one of each class per run), as a result of averaging images of the same class. Although there is not a fixed number of examples necessary to train the classifier, the more the better. Hebart et al. (2016) showed that run-wise beta estimates can be more accurate than single-trial ones, which can potentially lead to higher accuracies (Ku et al., 2008) or slightly improve power (Allefeld and Haynes, 2014). However, according to Pereira et al. (2008), at least a few tens of examples in each class are needed to properly estimate the parameters of the classifier, so LSS or LSA would be the most recommended option.

When trying to compare different methodological alternatives for decoding analysis, measuring the performance of the classifier is important, but evaluating its significance is crucial. In neuroscience research, the main aim is to determine the probability of a decoding result at the group level. The large number of voxels in fMRI analyses results in massive statistical tests, which need to be corrected for multiple comparisons. Cluster-level inference has become the most popular method due to its larger sensitivity compared to voxel-level inference. As the name suggests, this method does not estimate the false-positive probability of isolated voxels, but evaluates if a cluster is significant as a whole. To do so, this approach assumes that there is a correlation between adjacent voxels, so that the signal in each voxel is not completely independent of its neighbors. Cluster-level inference consists of two stages: first, a primary threshold at the voxel level is employed to obtain those that surpass a certain statistical *p*-value. The election of the threshold is arbitrary in some way (Friston et al., 1994), and what is more important, results can highly vary depending on the threshold considered. Setting a liberal primary threshold may decrease the spatial specificity, in addition to boosting the false-positives rate. In fact, Woo and Wager (2014) demonstrated that using primary thresholds that are too liberal can have detrimental effects on false positives, localization and interpretation. Regarding the second stage, a cluster-level extent threshold is used to retain the set of voxels that surpass the minimum size that a cluster should have to be considered significant. This threshold is computed based on theoretical methods such as Random Field Theory (RFT, Worsley et al., 1999), Monte Carlo simulations (Forman et al., 1995) or non-parametric approaches (Nichols and Holmes, 2002).

Previous studies have shown that RFT corrections tend to be too conservative as well as prone to false positives (Eklund et al., 2016). Besides, RFT carries several assumptions about the data which are not always met, such as the smoothness of the fMRI images or the uniform distribution of this smoothness over the brain. However, the key problem for applying RFT in classification-based analysis is that the distribution of the accuracies is unknown, and they are assumed to be normally distributed. As an alternative, statistical significance can be evaluated by non-parametric approaches based on permutation testing, which does not require any assumption except exchangeability. The basic principles of permutation testing are simple (Brammer et al., 1997; Bullmore et al., 1999; Chen et al., 2011; Nichols and Holmes, 2002; Pereira and Botvinick, 2011; Winkler et al., 2014), and previous research has theoretically evaluated their use in classification analyses (Golland and Fischl, 2003). Briefly, this test consists on shuffling the data, computing statistics and cluster sizes and generating a null distribution of the cluster sizes, from which it is possible to establish the minimum size needed to reach significance (see Nichols and Holmes (2001) for a more detailed explanation). Based on this concept, Stelzer et al. (2013) proposed a framework o derive a cluster size *p*-value at the group level, employing a Monte Carlo method to combine individual results. To compute the cluster-defining primary threshold, this method builds an empirical distribution for each voxel separately, minimizing the consequences related to spatial inhomogeneities that a global accuracy threshold would have. An alternative solution was proposed by Smith and Nichols (2009), the so-called Threshold-Free Cluster Enhancement (TFCE). This algorithm transforms the value of each voxel to a weighted score of the surrounding voxels, summarizing the cluster-wise evidence at each voxel. However, its most interesting contribution is that it does not require setting a cluster-defining primary threshold, eliminating the arbitrariness on this election and the subsequent impact on the results.

Previous research has compared how different pattern estimation methods compute the activity elicited by each trial separately. However, frequently, paradigms aim at isolating the activity of different events within the same trial, which suffers from significantly high signal overlap. The effect of alternative methods in this type of experimental design is yet unknown. Therefore, in this study, we aimed at evaluating the performance of different approaches in a context where a sustained activity had to be isolated from a zero-duration event (Dataset 1). Specifically, we tested the performance of LSU, LSA and LSS methods in the aforementioned design (Dataset 1), in addition to a classic block design (Dataset 2) and an event-related design (Dataset 3). Based on previous studies (Abdulrahman and Henson, 2016; Mumford et al., 2014, 2012), we predicted that LSS would estimate more accurately the signal elicited by each trial event, due to the way this method addresses the collinearity between closein-time experimental conditions. This collinearity is lower both in blocked or slow event-related designs, so that the three pattern estimation methods should be able to accurately estimate the activation patterns. Moreover, we examined the suitability of parametric (*t*-test) and non-parametric (Stelzer’s and TFCE) approaches to evaluate the significance of the results obtained with the different estimation methods. We hypothesized that the two non-parametric techniques would yield a higher sensitivity than the standard *t*-test, although variations between the two permutation-based approaches were expected due to the different cluster-search algorithms that they employ and the way permutations are applied. In contrast, we predicted that the three pattern estimation methods and the different statistical approaches would obtain similar results in the block design.

## 2. Material and Methods

### 2.1. Dataset 1

#### 2.1.1. Participants

Twenty-four students from the University of Granada (M = 21.08, SD = 2.92, 12 men) took part in the experiment and received an economic remuneration (20-25 euros, according to performance). All of them were right-handed with normal to corrected-to-normal vision, no history of neurological disorders, and signed a consent form approved by the local Ethics Committee.

#### 2.1.2. Acquisition

fMRI data were acquired using a 3T Siemens Trio scanner at the Mind, Brain and Behavior Research Centre (CIMCYC) in Granada (Spain). Functional images were obtained with a T2*-weighted echo planar imaging (EPI) sequence, with a TR of 2000 ms. Thirty-two descendent slices with a thickness of 3.5 mm (20% gap) were obtained (TE = 30 ms, flip angle = 80°, voxel size of 3.5 mm^3^). The sequence was divided in 8 runs, consisting of 166 volumes each. After the functional sessions, a structural image of each participant with a high-resolution T1-weighted sequence (TR = 1900 ms; TE = 2.52 ms; flip angle = 9°, voxel size of 1 mm^3^) was acquired.

We used SPM12 (http://www.fil.ion.ucl.ac.uk/spm/software/spm12) to preprocess and analyze the neuroimaging data. The first 3 volumes were discarded to allow for saturation of the signal. Images were realigned and unwarped to correct for head motion, followed by slice-timing correction. Afterwards, T1 images were coregistered with the realigned functional images. To better preserve the spatial configuration of activations in individual subjects, images were not smoothed nor spatially normalized into a common space.

#### 2.1.3. Design

The task contained two events in each trial, first a word (positive, negative or neutral in valence) and second two numbers, to which participants had to respond. These two numbers corresponded to the offer that participants received, from which they decided to accept or not based on the fairness/unfairness of the offer. They performed a total of 192 trials, arranged in 8 runs (24 trials per run), in a counterbalanced order across participants. Each trial started with the word for 1000 ms, followed by a jittered interval lasting 5500 ms on average (4-7 s, +/0.25°). Then, the numbers appeared for 500 ms followed by a second jittered interval (5500 ms on average; 4-7 s, +/0.25°). The first event was modelled as the duration of the word and the variable jittered interval, yielding a global duration ranging from 5 to 8 seconds. The second event was modelled as an impulse function (Dirac delta), i.e. with zero duration, as explained in Henson (2005). Participants read an adjective with a certain valence, and then they used this information to prepare to respond to the offer (second event). Thus, there is a preparatory process that leads to a sustained activity along time. However, the second event captures a completely different process. Once participants make a decision (accept or not), the process ends. A large body of literature shows that preparatory processes extend in time (e.g. Bode and Haynes, 2009; González-García et al., 2017, 2016; Sakai, 2008) whereas responding to a brief target does not (see the temporal duration of the potentials in Moser et al., 2014). This has been also measured by other neuroimaging methods, such as the CNV ERP potential (Russo et al., 2017). For this theoretical cognitive reason, this second event was modelled with zero duration. Besides, we ran an additional analysis to evaluate whether modelling the first event as an impulse function (zero duration) influenced the results by reducing the collinearity of the regressors between the first and second events in a trial. On the other hand, the beginning of runs and the inter-trial jittered intervals served as the implicit baseline. The whole fMRI session lasted 41 minutes approximately.

To test the performance of the different approaches (accurate estimation of signal activity for pattern estimation methods and large sensitivity and low false-positives rate for statistical methods), we focused on two different classification analyses, one for each part of the trial. We firstly aimed at discriminating the positive *vs.* negative valence of the words (e.g. Lindquist et al., 2015; from now on, *valence* classification) that were equated in number of letters, frequency of use and arousal (Gaertig et al., 2012). The total number of images available for the classification procedure varied according to the method used to estimate the patterns. As Table 1 shows, LSU yielded 8 images per condition, one for each run. LSA and LSS obtained the same number as positive/negative trials in the experiment (64 of each category, per participant). Last, we aimed to discriminate between fair and unfair offers (*fairness* classification). LSU yielded again 8 images per condition. On the other hand, LSA and LSS obtained 96 images for each condition and participant.

**Table 1:** Average number of beta maps obtained by each pattern estimation method and dataset used, for each classification problem evaluated

|  | Dataset 1 |  | Dataset 2 | Dataset 3 |
| --- | --- | --- | --- | --- |
| Method | Valence | Fairness | Faces vs Houses | Familiarity |
| LSU | 8 | 8 | 12 | 11 |
| LSA | 64 | 96 | 12 | 176 |
| LSS | 64 | 96 | 12 | 176 |

**Table 2:**
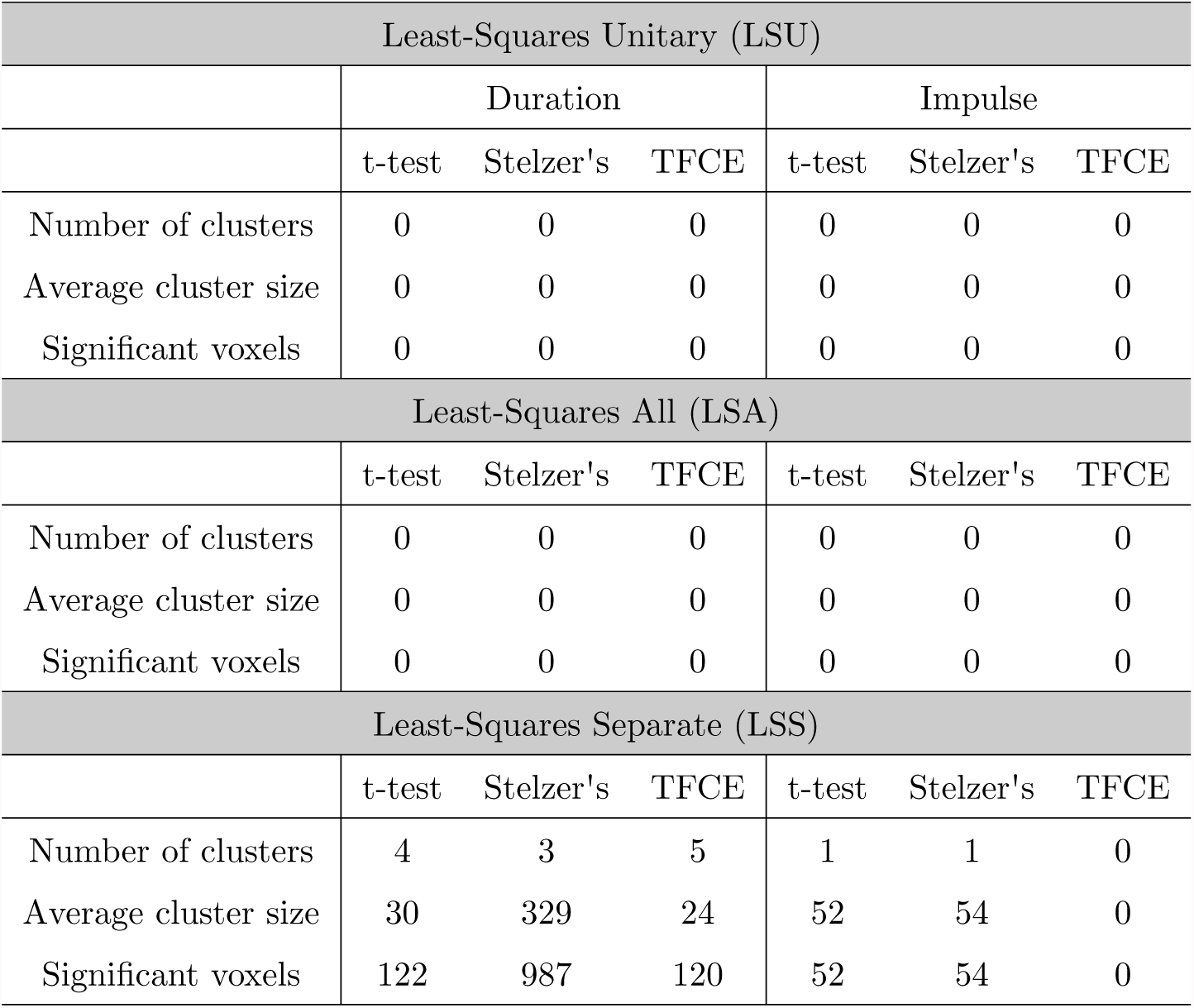
Comparison of the clusters distribution by the different pattern estimation methods and statistical tests in the *valence* classification after modelling the words as epochs/zeroduration events.

### 2.2. Dataset 2

We used data of six participants from the study published by Haxby et al. (2001), which has served as example fMRI dataset several times (e.g. Hanson et al., 2004; O’Toole et al., 2007). Neural responses were measured with gradient echoplanar imaging on a GE 3T scanner (General Electric, Milwaukee, WI) [repetition time (TR) = 2500 ms, 40 3.5-mm-thick sagittal images, field of view (FOV) = 24 cm, echo time (TE) = 30 ms, flip angle = 90°] while they performed a one-back repetition detection task. High-resolution T1-weighted spoiled gradient recall (SPGR) images were obtained for each subject to provide detailed anatomy (124 1.2-mm-thick sagittal images, FOV = 24 cm).

The dataset consists of 12 runs where the participants viewed grayscale images of eight object categories: faces, houses, cats, bottles, scissors, shoes, chairs and scrambled images. Each run began and ended with 12-s rest and contained eight blocks of 24-s duration, one for each category, separated by 12- s of rest. Stimuli were presented for 500 ms with an interstimulus interval of 1500 ms. We focused on the faces *vs.* houses classification, although the rest of the stimuli were also included in the GLM to preserve the implicit baseline. Since only one block for each stimulus type was presented in each run, LSU and LSA were equivalent. Although the LSS estimation was developed for event-related designs, we implemented a blocked-version of the LSS approach by iteratively fitting a new GLM for each block. For each model, the target condition is associated with one regressor, and the rest are associated with one error regressor. Thus, there are 8 models for each run, one for category. All methods yielded the same number of estimates to train the algorithm: 1 per run and condition.

### 2.3. Dataset 3

We used data from 33 participants of a recent study published by Visconti di Oleggio Castello et al. (2017). The full database was openly available in Datalad repository (http://datalad.org). Brain images were acquired using a 3T Philips Achieva Intera scanner with a 32-channel head coil [repetition time (TR) = 2000 ms, 35 3-mm-thick axial images, field of view (FOV) = 24 cm, echo time (TE) = 35 ms, flip angle = 90°]. A single high-resolution T1-weighted (TE/TR = 3.7/8.2 ms) anatomical scan was acquired with a 3D-TFE sequence. For a more detailed explanation see the original work (Visconti di Oleggio Castello et al., 2017). Preprocessing was carried out following the same procedure used for Dataset 1.

The dataset consists of 11 runs where the participants viewed pictures portraying different familiar and unfamiliar identities: four faces of friends, four unknown faces, and the participant’s own face. A trial consisted of three different images of the same individual (normal trial) or two different identities (odball trials), each presented for 500 ms with no gap, followed by a 4500 ms inter-trial interval displaying a white fixation cross. Each trial was modelled with a duration of 1.5 seconds, as it was done in the original paper (Visconti di Oleggio Castello et al., 2017). The order of the events was pseudo-randomized to approximate a first-order counterbalancing of conditions. A functional run contained 48 trials: four trials for each of the nine individuals (four familiar, four unfamiliar and self), four blank trials, four oddball and four buffer trials (three at the beginning and one at the end). Each run had 10 seconds of fixation at the beginning to stabilize the BOLD signal and at the end (to collect the response to the last trials). We focused on discriminating the neural activity associated with familiar *vs.* unfamiliar faces although the rest of the stimuli were also included in the GLM to preserve the implicit baseline. Eleven beta estimates per condition were obtained by LSU, whereas LSA and LSS yielded 176.

### 2.4. Searchlight analysis

For each dataset, we employed a searchlight approach across the whole brain (Kriegeskorte et al., 2006). We used The Decoding Toolbox (TDT, Hebart et al., 2015) to create spherical regions of 12 mm, limiting the analysis to the voxels contained in it. This size was chosen according to previous studies that showed a systematic decrease in performance when a larger size is selected (e.g. Arco et al., 2016; Chen et al., 2011). The procedure was repeated across all the positions of the brain, yielding an accuracy map in which each value represented the accuracy obtained when a given voxel was the center of the sphere. To classify images, we employed a support vector machine (SVM) with a linear kernel (Misaki et al., 2010; Pereira et al., 2008). A leave-one-run-out scheme was used to cross-validate the performance of the classifier (Coutanche and Thompson-Schill, 2012; Haynes and Rees, 2006; Lee et al., 2011; Reddy et al., 2010; Wolbers et al., 2011). In this scheme, the classifier is trained with images from all but one run, whereas the patterns of the remaining run are used to test the performance of the algorithm. The number of images available for the training/testing process highly depends on the dataset used, the pattern estimation method employed and the classification problem evaluated. This information is summarized in Table 1.

### 2.5. Evaluating statistical significance

The use of multivariate decoding for interpretation instead of prediction does not aim at obtaining a classifier with the largest accuracy as possible, but obtaining a decoding model that performs reliably better than chance (Hebart and Baker, 2017). This would demonstrate that there is information in the data related to the experimental condition under study, which increases our knowledge about the neural mechanisms associated with a certain cognitive function. Moreover, there is a certain variability between each individual brain, so it is necessary to evaluate if the obtained results are significant at the population level. We describe in this section the theoretical framework of the three methods employed in our study.

#### 2.5.1. t-test and Gaussian Random Field Theory

The first method evaluated is based on Gaussian Random Field Theory (RFT), a mathematical framework that finds the specific threshold for a smooth statistical map that meets the required family-wise error rate (Brett et al., 2003). The smoothness of a statistical image is not usually known, but it can be estimated as the number of resels that the image has. The concept of resel (resolution element), introduced in Worsley et al. (1992), is similar to the number of independent observations in the image, and it is a function of the number of voxels in the image and the Full Width at Half Maximum (FWHM). Another crucial concept is the Euler characteristic (EC), which is a property related to the probability that a number of clusters is considered significant when a certain statistical threshold is used. Following the expression derived from Worsley et al. (1992), it is possible to compute the expected value of EC, as follows:
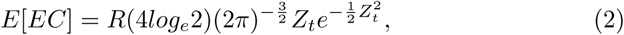

where *R* is the number of resels and *Z_t_* is the *Z* score threshold. This expression corresponds to images of two dimensions, but the methods are equivalent to three-dimensional images. It is worth noting that the expected value of the EC is approximately equivalent to the probability of a family wise error, especially at high thresholds. By setting this value to the standard 0.05, it is possible to conclude that the remaining clusters have a maximum probability of 0.05 of having occurred by chance.

We employed the functions provided by the SPM12 package (http://www.fil.ion.ucl.ac.uk/spm/software/spm12) to apply this method, modifying a line of code to take into account the non-stationarity of the underlying random field. The procedure followed was the same for all the datasets evaluated. After computing the decoding accuracy map for each subject, all maps were normalized to a standard EPI. Then, a voxel-wise *t*-test against the theoretical chance (0.5 in our binary-classification analyses) was applied to these normalized maps. We employed a cluster-defining primary threshold of *p <* 0.001 (uncorrected), which was later used to find significant clusters (FWE corrected, *p <* 0.05) on the resulting map.

#### 2.5.2. Stelzer’s

The second method evaluated, Stelzer’s, combines results from each subject with a Monte Carlo method and based on that, derives a cluster size *p*-value at the group level. This approach is based on permutation tests, which unlike RFT, rely on minimal assumptions. Specifically, a within-subject searchlight analysis was performed shuffling the labels corresponding to the two experimental conditions to distinguish from. We carried out this step 100 times per participant, yielding 100 permuted accuracy maps. Then, these maps were spatially-normalized to a standard EPI image to register images of different subjects into the same coordinate system. A map from each participant was randomly picked following a Monte Carlo resampling with replacement (Forman et al., 1995), averaging the values voxel-wise and obtaining a permuted group map. This procedure was carried out 50000 times, yielding 50000 group permuted maps. This process is equivalent to building an empirical chance distribution for each voxel in the brain. To evaluate the significance of each voxel, it is necessary to compare the null distribution with the real accuracy. This accuracy is obtained by training the classifier with actual true labels, and averaging the resulting maps across subjects (from now on, the real group map). For a cluster-defining primary threshold of *p*-value = 0.001 and a distribution of 50000 samples, a voxel will be significant when no more than 50 voxels of the empirical distribution have a larger value than the value of the real group map. To compute the specific *p*-value for a voxel *x,* we employed the following equation:
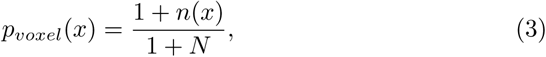

where *n*(*x*) is the number of samples from the empirical distribution with a larger value than the one obtained training the classifier with the true labels at the voxel *x,* and *N* is the number of permutations done.

Once the image has been thresholded at the voxel-level (applying the cluster-defining primary threshold), an empirical distribution of the cluster sizes of the 50000 permuted maps is built to compute the required family-wise error rate at the cluster-level. A set of contiguous voxels are considered a cluster if they share a face, but not an edge or a vertex, in which Stelzer et al. (2013) defines as a 6-connectivity scheme. This cluster search is also applied to the real group map, so that only the clusters which surpass the cluster-level extent threshold are considered significant. A cluster with a size *s* is computed to have a *p*-value of
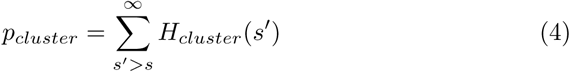

where *H_cluster_* is the normalized histogram of cluster sizes in the empirical distribution (number of clusters with size *s’* divided by the total number of clusters). Once each cluster size has an associated *p*-value, an FWE correction (*p* = 0.05) is applied on all clusters *p*-values to correct for multiple comparisons at the cluster level. The whole procedure is summarized in Figure 4.

**Figure 4:**
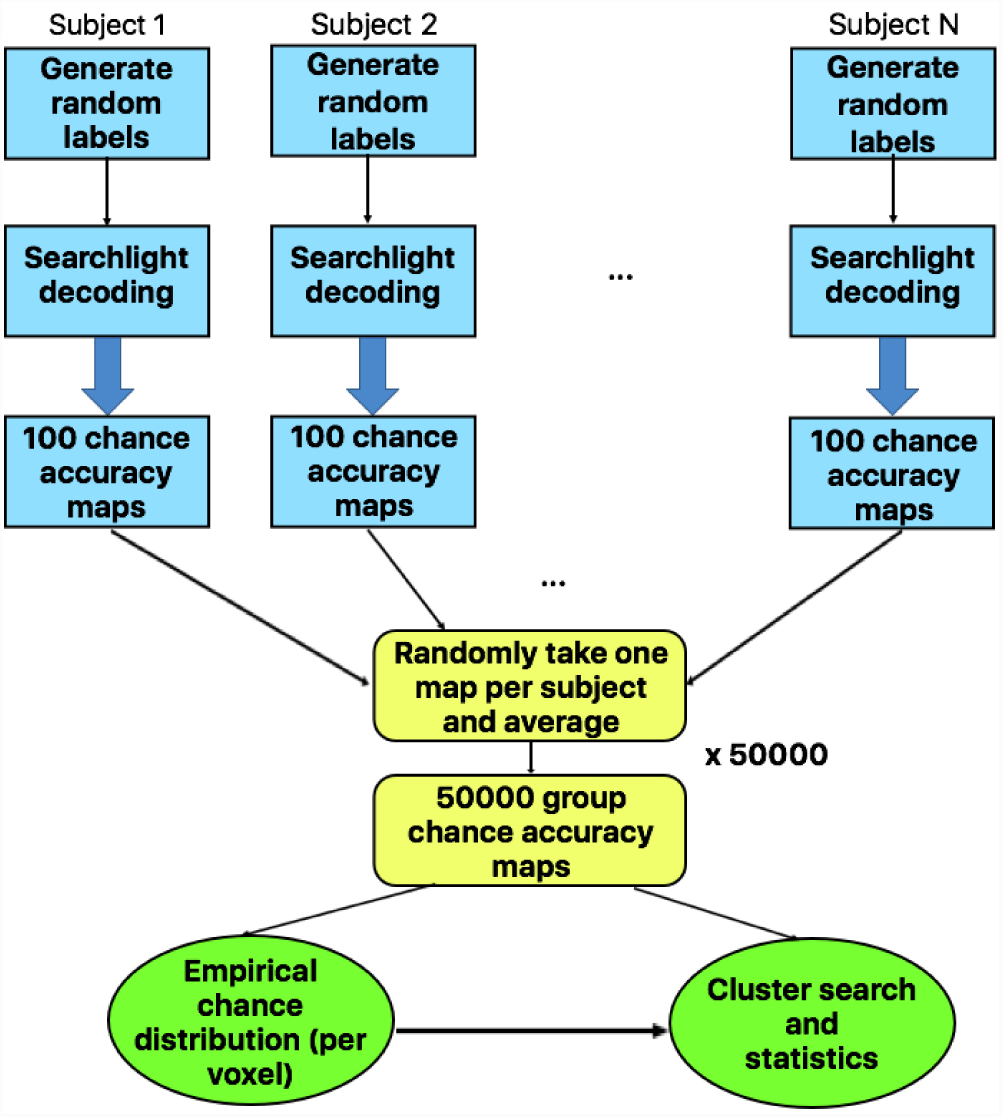
Schematic representation of Stelzer’s method. For each subject, a classifier is trained 100 times permuting the labels of the images, resulting in 100 accuracy maps which are spatially normalized into a common space. From each subject, a map is randomly picked following a Monte Carlo resampling with replacement procedure, averaging the values voxelwise to obtain a permuted group map. This procedure is repeated 50000 times, building empirically a chance distribution for each voxel position and selecting the 50th greatest value, which statistically corresponds to the accuracy threshold that marks the significance.

Figure 5 shows an example of the group distribution of the accuracies in one voxel for Dataset 1 (*valence* classification) and Dataset 2. Training with permuted labels results in accuracies around chance level (50%) in most of the permutations. The green vertical line indicates the significance threshold at which a given accuracy is considered significant, whereas the black one shows the accuracy level obtained after training the classifier with the true labels. It is worth noting that accuracies are not homogeneous across the brain, but they depend on the region from which information is being decoded. For this reason, it is remarkable that this method computes a different empirical distribution for each voxel separately. We employed custom code to carry out all the described stages of Stelzer’s method.

**Figure 5:**
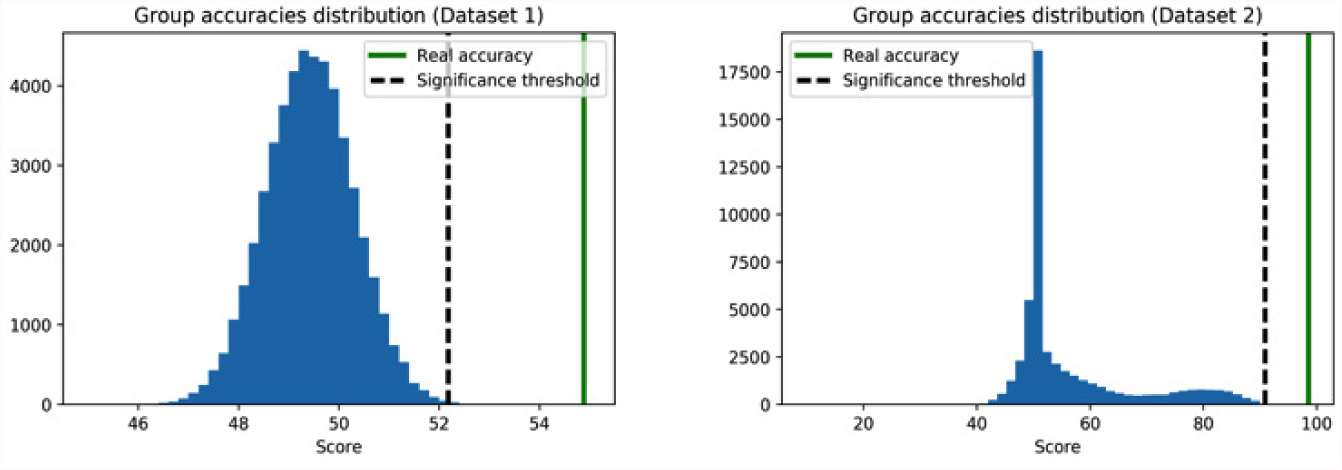
Distributions of group permuted accuracies in one voxel for the two datasets used: Dataset 1 (left) and Dataset 2 (right). While in Dataset 1 most accuracies are around chance level, in the second one the number of voxels that surpass the threshold is much larger.

#### 2.5.3. TFCE

The last method used was TFCE, included in the FMRIB Software Library (FSL; https://fsl.fmrib.ox.ac.uk/fsl/fslwiki). The basis of this method is to transform images to facilitate the discrimination between significant and non significant voxels. This transformation relies on the concept that in each image, there are sets of contiguous voxels which are candidates to belong to a cluster. There are two possible extreme scenarios: the first one is that the intensity of the voxels is large (high statistical values) but they are locally distributed. However, it is also possible that the signal is weak (low statistical values) but spatially extended. The main aim of TFCE is to level these two situations so that it is equally likely that both cases reach significance. Mathematically, the expression to compute a TFCE score is
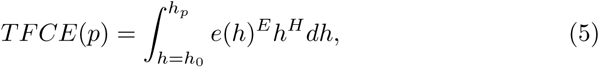

where *h_o_* is typically zero, *e* is the extent of the cluster that voxel *p* belongs to and *h* is the primary threshold. For each voxel, a TFCE score is computed as the sum of the product between the extent of the cluster and the different primary thresholds (*h* ranging from 0 to *h_p_*). The contribution of these two factors depends on *E* and *H.* Smith and Nichols (2009) evaluated a wide range of values for these parameters and established that *E* = 0.5 and *H* = 2 are the optimal ones.

In our analysis following this approach, the accuracy maps for all participants were entered into a second-level analysis, where a one-sample *t*-test was used to contrast conditions. To assess significance at the population level, permutation tests were applied. On each permutation, the signs of the individual accuracy maps were randomly flipped and a new *t*-test was performed. This was repeated 50000 times, obtaining an empirical null distribution of t-values. The TFCE transformation was later applied to find significant clusters (FWE-corrected, *p* = 0.05). Figure 6 illustrates this procedure.

**Figure 6:**
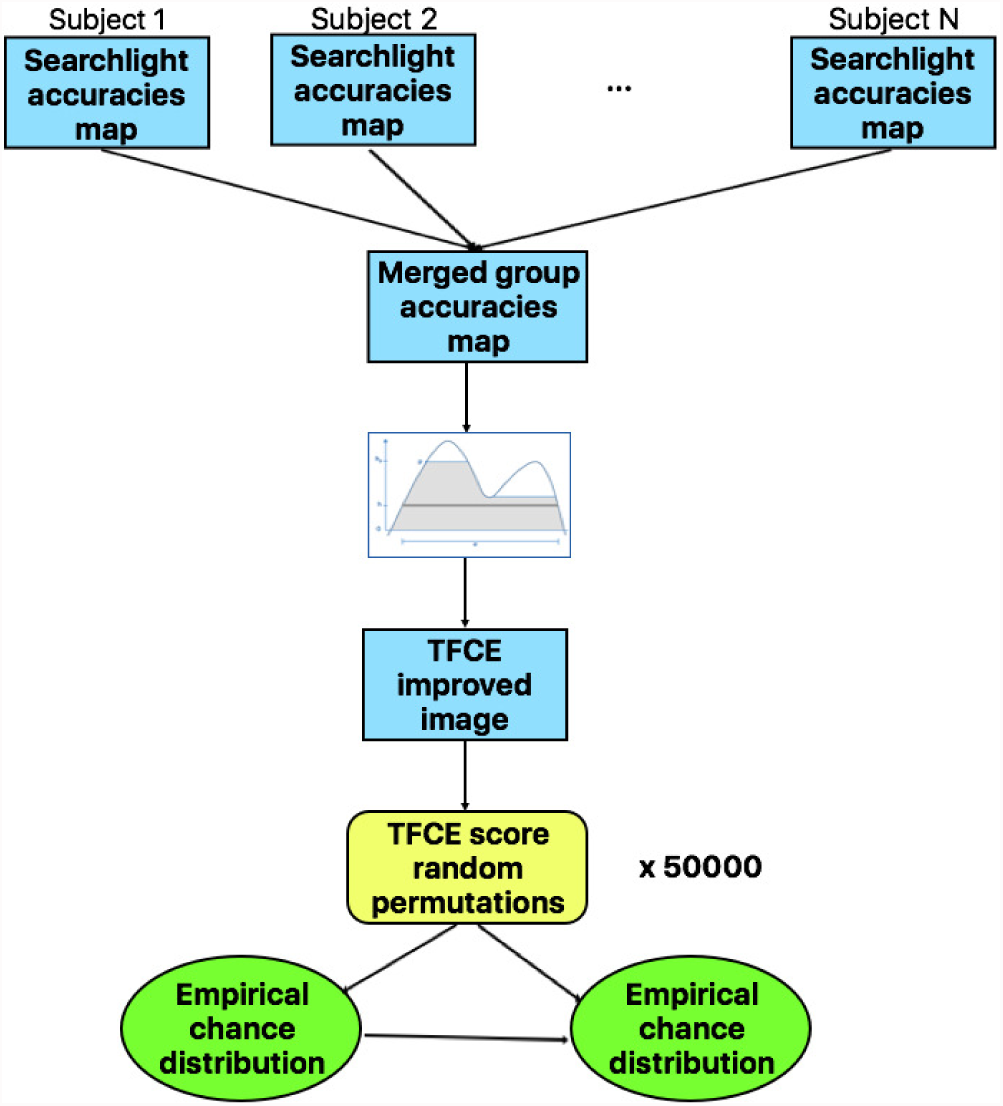
Schematic representation of the TFCE approach. Once all subjects’ accuracy maps are merged, the TFCE algorithm is applied. For a given point *p,* its value is replaced by an average of the intensities of the voxels of its neighbourhood, enhancing the intensity within cluster-like regions. To determine the significant voxels, a one-sample *t*-test is used. Besides, permutation tests are applied, flipping the signs of the individual accuracy maps and performing a new *t*-test. To correct for multiple comparisons, the null distribution of the maximum TFCE score is built up, testing the actual TFCE image against it.

## 3. Results

In this section, we report the results from the three datasets evaluated in this study (1: two events of different duration in each trial, 2: block design, 3: event related design with events of the same non-zero duration) estimated with LSU, LSA and LSS and statistically tested with parametric (*t*-test) and non-parametric (Stelzer’s and TFCE) approaches. Additionally, we evaluated two different ways of modelling the two events in Dataset 1 to study how the duration of the events influenced the results. In the first one, the duration of the jittered interval that separates the two events was added to the first event (words). The alternative was to model both events (words and numbers) as impulse functions, i.e. with zero duration.

### 3.1. Comparison of different pattern estimation models (LSU, LSA and LSS)

We first focused on comparing the three pattern estimation methods in four different scenarios: *i*) a paradigm with two events of different durations per trial (event-related design) where the individual contribution of both events was computed, Dataset 1,*ii*) same paradigm but modelling the two events with zero duration, Dataset 1, *iii)* a block-design from the pioneering study of Haxby et al. (2001), Dataset 2, and *iv)* an event-related design from a recently published study (Visconti di Oleggio Castello et al., 2017) where all trials were modelled with the same duration, Dataset 3.

Results in terms of cluster detection and number of significant voxels observed are summarized in Tables from 2 to 4. For the *valence* classification in Dataset 1, no significant voxels were found when LSU or LSA were applied regardless of the statistical method used and the way that events were modelled, whereas the LSS method uncovered a set of informative regions (see Figure 7 and 10). In the *fairness* classification, LSA was the only method that did not obtain any significant result. Table 3 summarizes the results for the *fairness* classification, illustrating very similar results for the two ways of modelling the first event. Besides, Figure 11 and Figure 13 show the small differences found between the two models. Regarding Dataset 2, all pattern estimation methods showed a larger sensitivity than with Dataset 1 (see Table 4). Specifically, the informative regions obtained by each one of them were very similar across methods. Unlike Dataset 1, LSA allowed a accurate estimation of the neural activity in Dataset 3, in addition to LSU and LSS. The experiment aimed at finding differences at the trial level and not to isolate the neural activity of different events within each trial, which is considerably harder.

**Table 3:** Comparison of the clusters distribution by the different pattern estimation methods and statistical tests in the *fairness* classification after modelling the words as epochs/zeroduration events.

| Least-Squares Unitary (LSU) |  |  |  |  |  |  |
| --- | --- | --- | --- | --- | --- | --- |
|  | Duration |  |  | Impulse |  |  |
|  | t-test | Stelzer's | TFCE | t-test | Stelzer's | TFCE |
| Number of clusters | 3 | 1 | 1 | 2 | 1 | 1 |
| Average cluster size | 628 | 15422 | 13909 | 1058 | 13832 | 14399 |
| Significant voxels | 1883 | 15422 | 13909 | 2116 | 13832 | 14399 |
| Least-Squares All (LSA) |  |  |  |  |  |  |
|  | t-test | Stelzer's | TFCE | t-test | Stelzer's | TFCE |
| Number of clusters | 0 | 0 | 0 | 0 | 0 | 0 |
| Average cluster size | 0 | 0 | 0 | 0 | 0 | 0 |
| Significant voxels | 0 | 0 | 0 | 0 | 0 | 0 |
| Least-Squares Separate (LSS) |  |  |  |  |  |  |
|  | t-test | Stelzer's | TFCE | t-test | Stelzer's | TFCE |
| Number of clusters | 2 | 1 | 1 | 1 | 1 | 1 |
| Average cluster size | 4469 | 17620 | 16790 | 9742 | 16342 | 13584 |
| Significant voxels | 8938 | 17620 | 16790 | 9742 | 16342 | 13584 |

**Figure 7:**
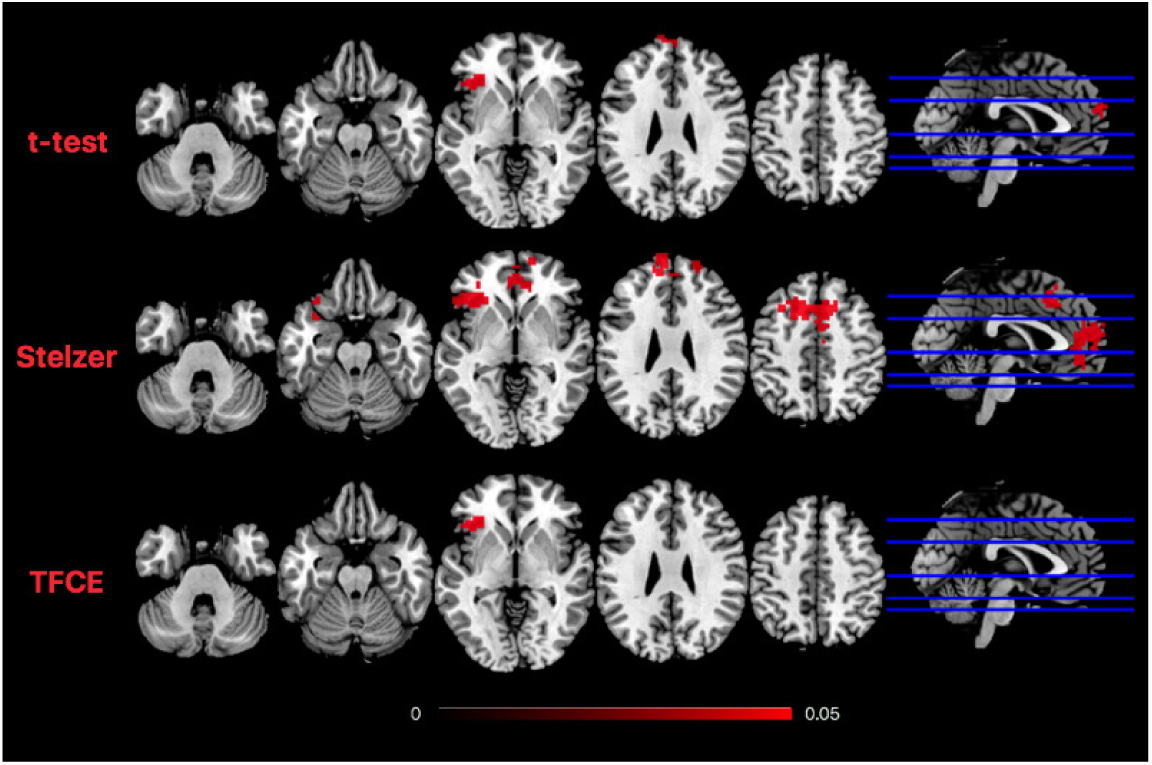
Significant results obtained by the LSS method when discriminating word valence in Database 1, modelling the words with its corresponding duration. Results from the *t*-test and TFCE are practically the same, both in location and number of significant voxels. Stelzer’s method, on the other hand, yields the significant regions obtained by the other methods while increasing the number of significant voxels, showing higher sensitivity.

**Figure 8:**
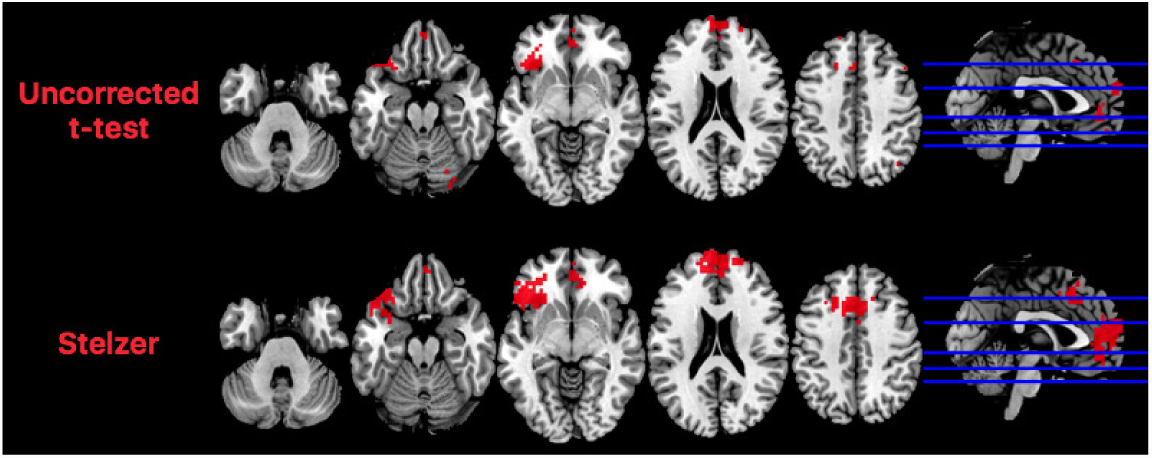
Comparison of the uncorrected results from the *t*-test (p G0.001) and the significant voxels obtained by Stelzer’s in Dataset 1 (modelling the duration of the words). The distribution of the voxels is similar in both cases, so that differences may rely on the inability to surpass the statistical threshold when the *t*-test is applied.

**Figure 9:**
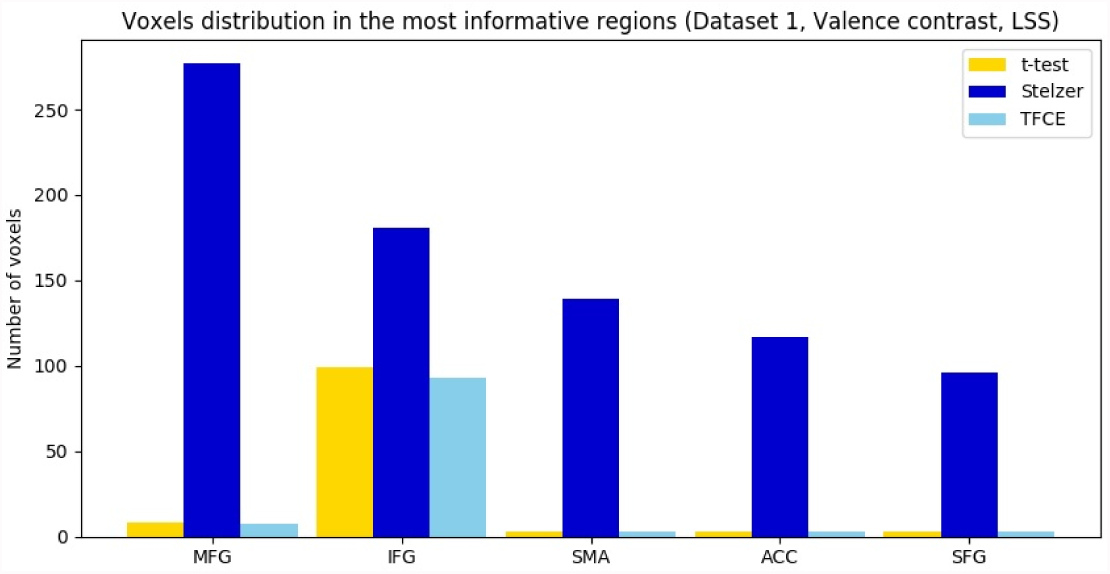
Voxel distribution in the most informative regions for the *valence* classification of Dataset 1, modelling the first event with its corresponding duration. The Inferior Frontal Gyrus is the only region where the three methods found informative voxels.

**Figure 10:**
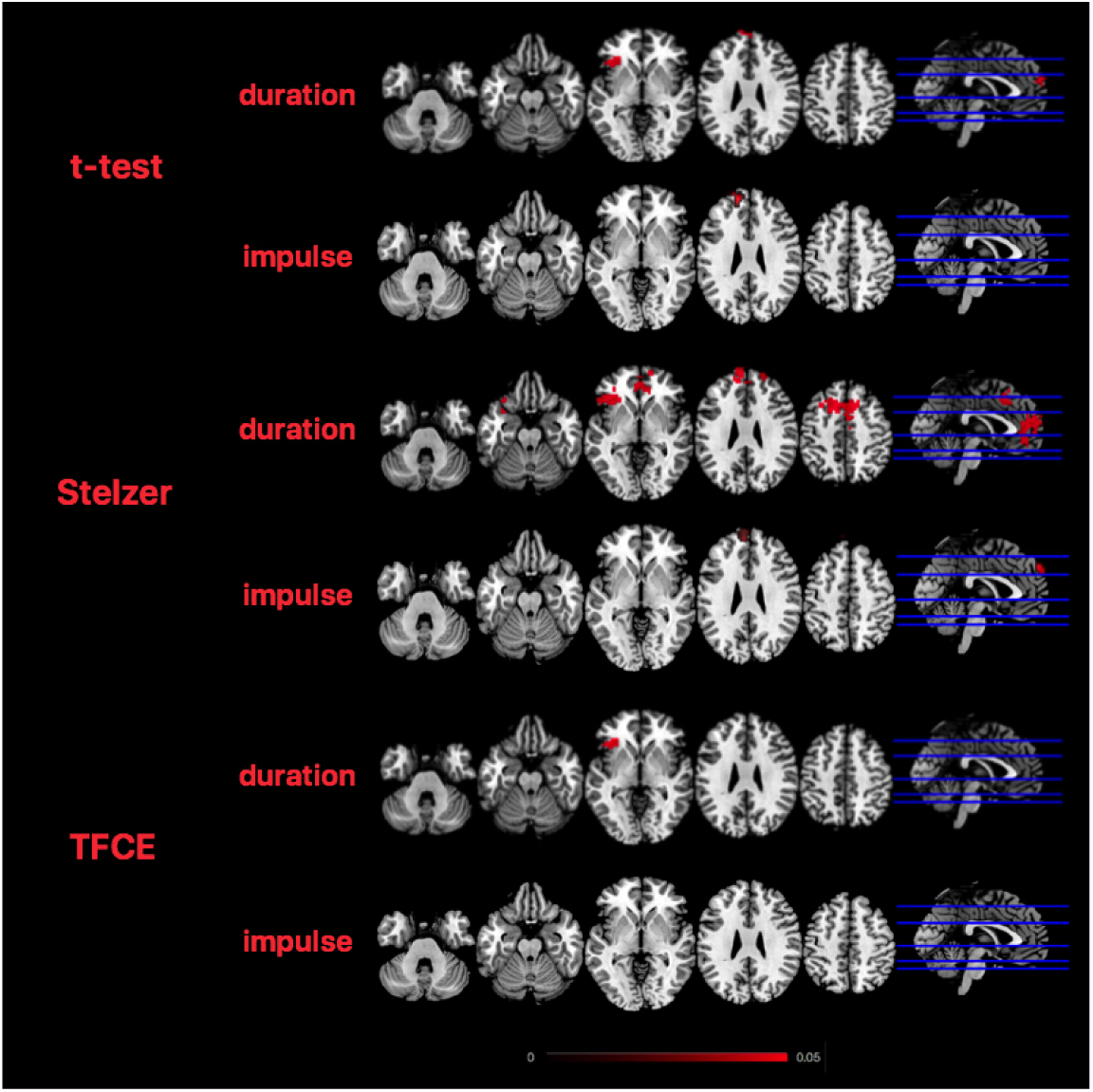
Comparison of the results obtained by the different pattern estimation and statistical methods in the *valence* classification modelling the words as duration/impulse events.

### 3.2. t-test vs. non-parametric methods

We next employed the three methods described in Section 2.5, that is, the *t-*test, Stelzer’s and TFCE, to assess significance of the obtained results. Figure 7 shows the significant results obtained by each of them when the LSS estimation method was employed in the *valence* classification of Dataset 1, when the first event was modelled with its corresponding duration. Here, the *t*-test and TFCE yielded essentially the same results in terms of number of voxels marked as significant and their spatial distribution, but largely differed from Stelzer’s. In fact, this method obtained approximately 8 times more significant voxels than the others. All clusters found by the *t*-test and TFCE were also included in Stelzer’s, but their spatial extent was larger in the latter. When the first event was modelled with zero duration, TFCE did not yield any significant result, but the *t*-test and Stelzer’s obtained exactly the same informative cluster. Figure 10 shows these results and illustrates the differences between the two ways of modelling the first event.

In the *fairness* classification (first event modelled with its corresponding duration), the differential sensitivity between the *t*-test and Stelzer’s was also obtained, but in this case, TFCE yielded very similar results to Stelzer’s instead than to the *t*-test (see Figure 11 and Figure 12). Modelling the first event as zero duration did not change much the results, as Figure 13 shows. It is important to highlight that when any of the non-parametric approaches was used, the difference in the informative regions obtained by the LSU and LSS methods was minimum. We further discuss the implications of this finding in Section 4.

**Figure 11:**
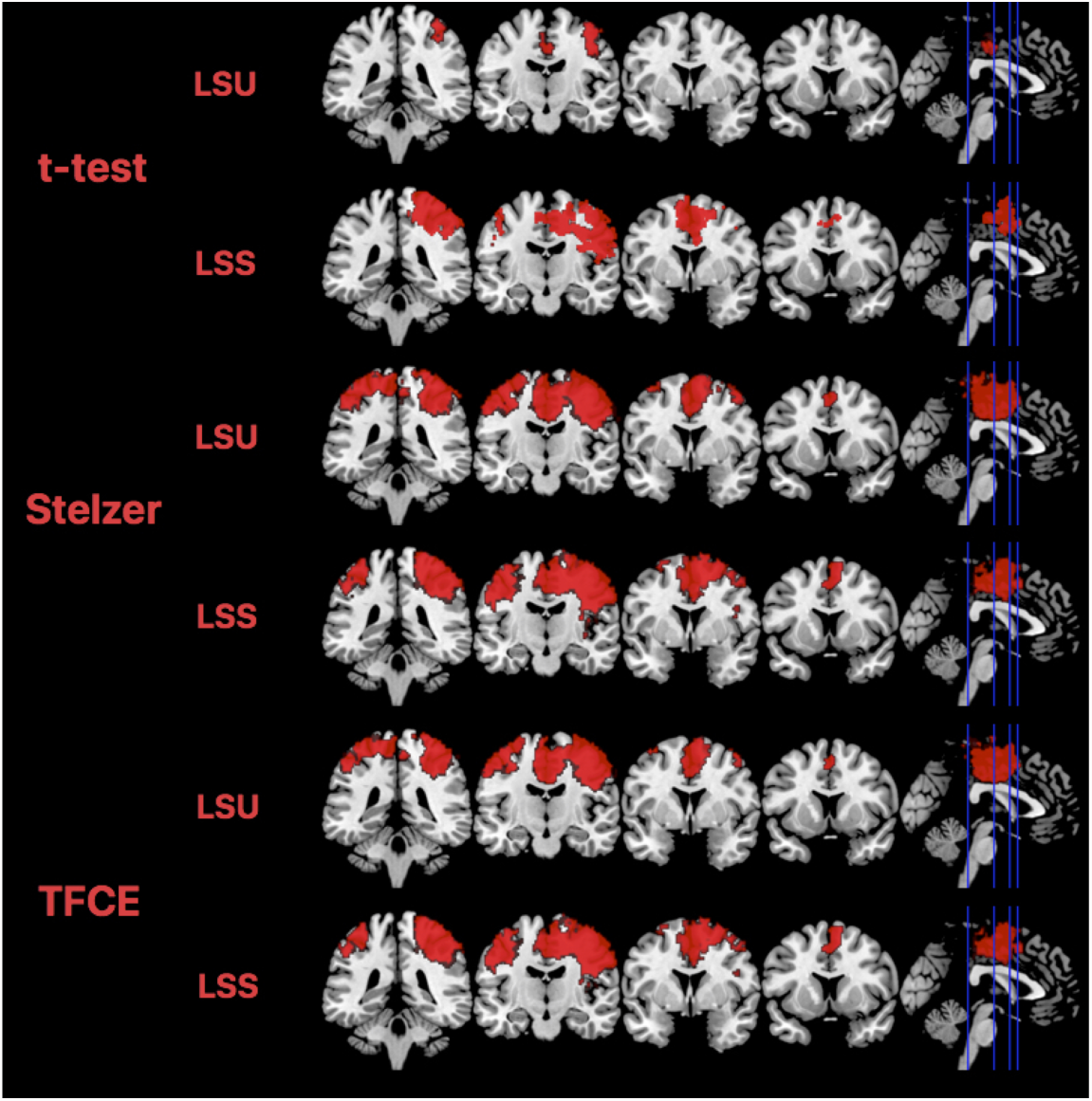
Significant results obtained by the different pattern estimation and statistical methods in the *fairness* classification modelling the words with its corresponding duration.

**Figure 12:**
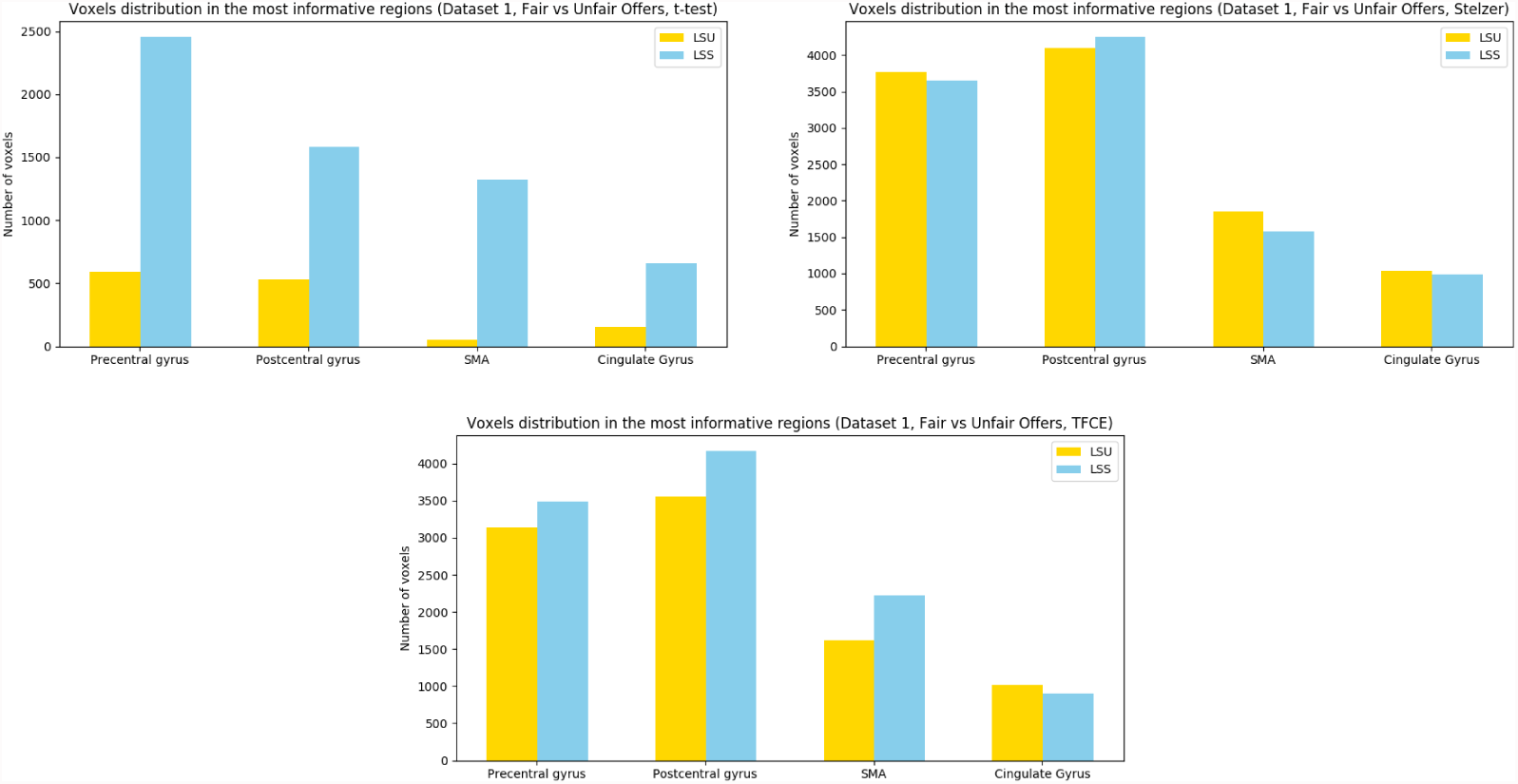
Voxel distribution in the most informative regions for the *fairness* classification of Dataset 1, modelling the first event with its corresponding duration. Region SMA = Supplementary Motor Area.

**Figure 13:**
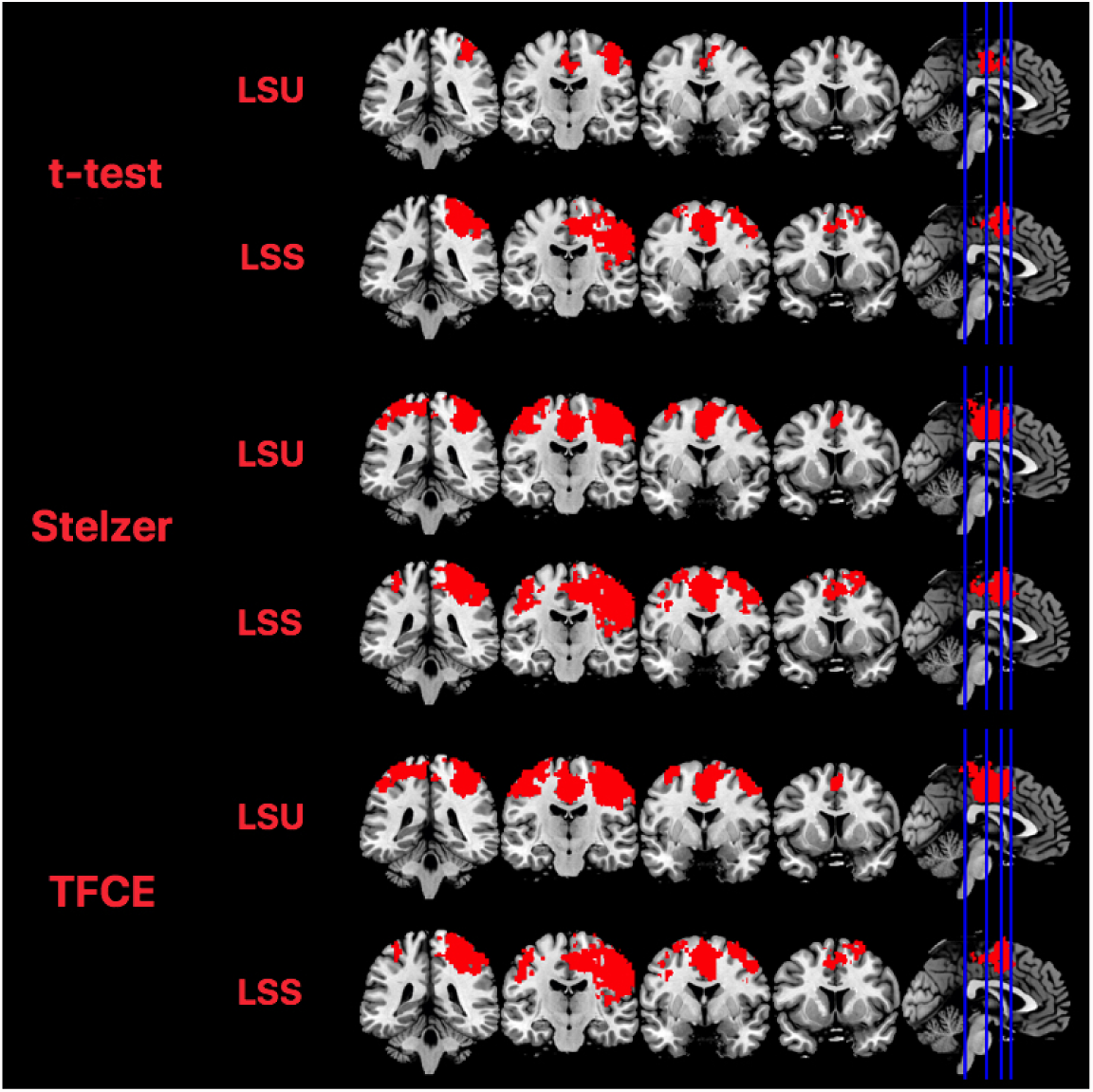
Significant results obtained by the different pattern estimation and statistical methods in the *fairness* classification of Dataset 1 where words were modelled as zero-duration events.

Figure 14 and Figure 15 reveal the differences between the three approaches for Dataset 2. Similarly to Dataset 1, Stelzer’s shows larger sensitivity regardless of the estimation method used, (see Table 4). Moreover, the location of the significant voxels is quite similar across the three approaches: they found a single massive significant cluster, slightly larger in case of TFCE and with a 35% of more significant voxels in the case of Stelzer’s in comparison with the *t*-test. This superior sensitivity of non-parametric methods is also observed in Dataset 3 (see Table 4), whereas the most informative brain regions are summarized in Figures 16 and 17.

**Figure 14:**
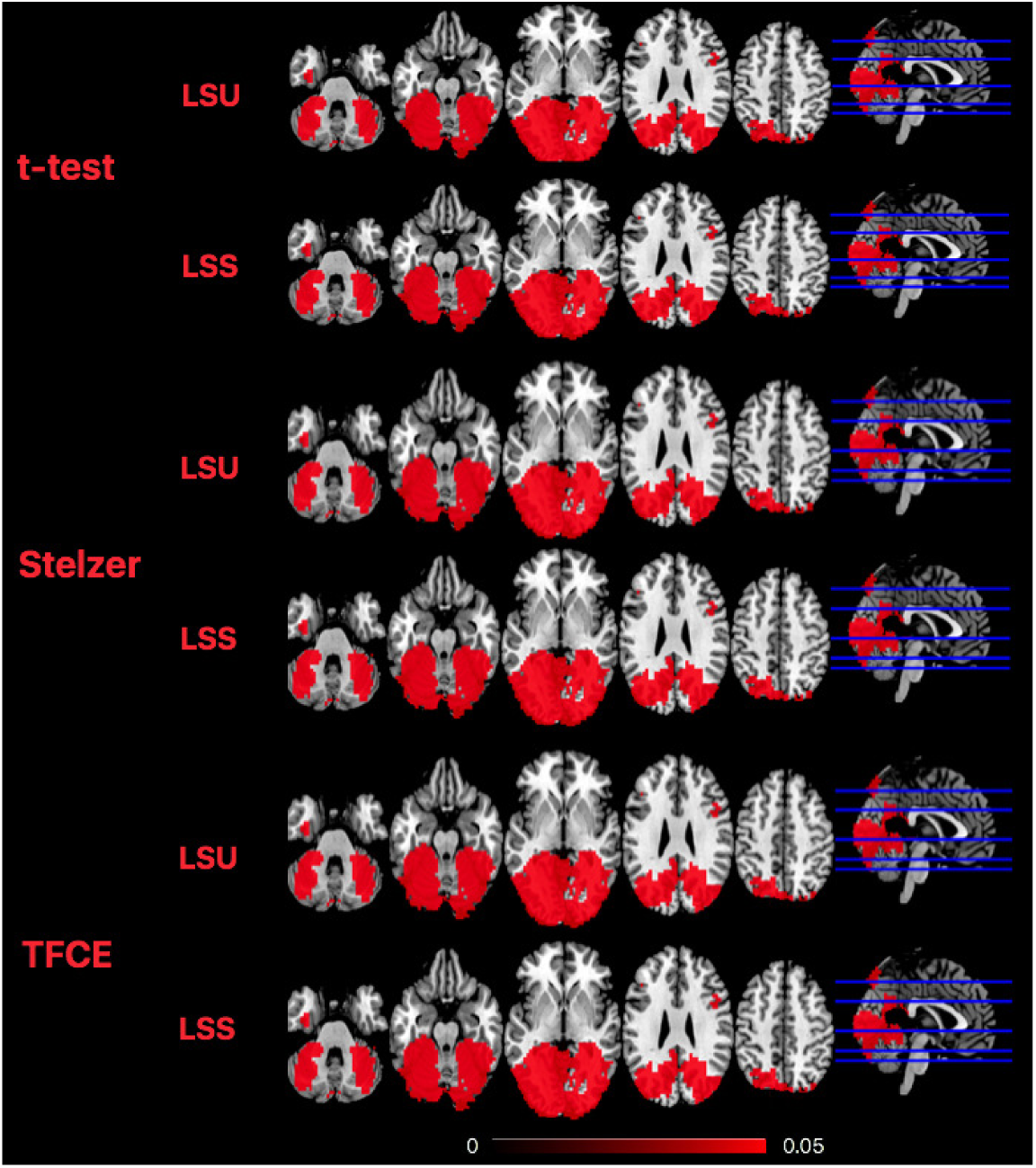
Significant results obtained by the different pattern estimation methods and techniques for evaluating the statistical significance in Haxby’s experiment. LSA is equivalent to LSU in this case, so only results for LSU and LSS are presented.

**Figure 15:**
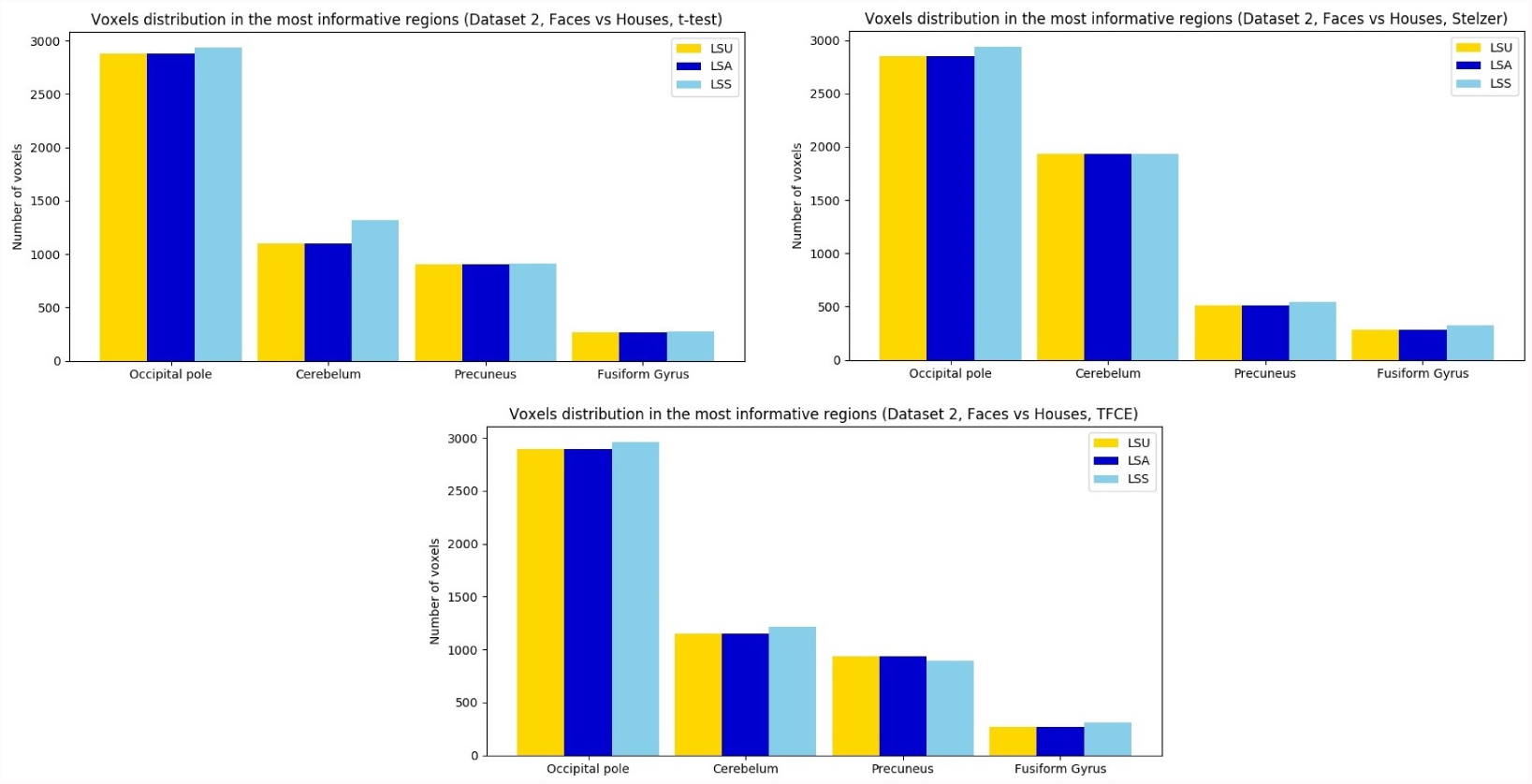
Voxel distribution in the most informative regions for each statistical and pattern estimation method in Dataset 2. Results are similar for all the methods employed, but discrepancy appears in the cerebelum and the precuneus.

**Table 4:** Summary of the clusters distribution by the different pattern estimation methods and statistical tests in Datasets 2 and 3.

| Least-Squares Unitary (LSU) |  |  |  |  |  |  |
| --- | --- | --- | --- | --- | --- | --- |
|  | Dataset 2 |  |  | Dataset 3 |  |  |
|  | t-test | Stelzer's | TFCE | t-test | Stelzer's | TFCE |
| Number of clusters | 4 | 1 | 1 | 5 | 2 | 3 |
| Average cluster size | 1821 | 9881 | 7717 | 527 | 2511 | 748 |
| Significant voxels | 7283 | 9881 | 7717 | 2635 | 5021 | 2244 |
| Least-Squares All (LSA) |  |  |  |  |  |  |
|  | t-test | Stelzer's | TFCE | t-test | Stelzer's | TFCE |
| Number of clusters | 4 | 1 | 1 | 2 | 2 | 1 |
| Average cluster size | 1821 | 9881 | 7717 | 1476 | 1383 | 4489 |
| Significant voxels | 7283 | 9881 | 7717 | 2952 | 2766 | 4489 |
| Least-Squares Separate (LSS) |  |  |  |  |  |  |
|  | t-test | Stelzer's | TFCE | t-test | Stelzer's | TFCE |
| Number of clusters | 4 | 1 | 1 | 2 | 3 | 1 |
| Average cluster size | 1831 | 9906 | 7692 | 1424 | 1463 | 4551 |
| Significant voxels | 7321 | 9906 | 7692 | 2847 | 4387 | 4551 |

**Figure 16:**
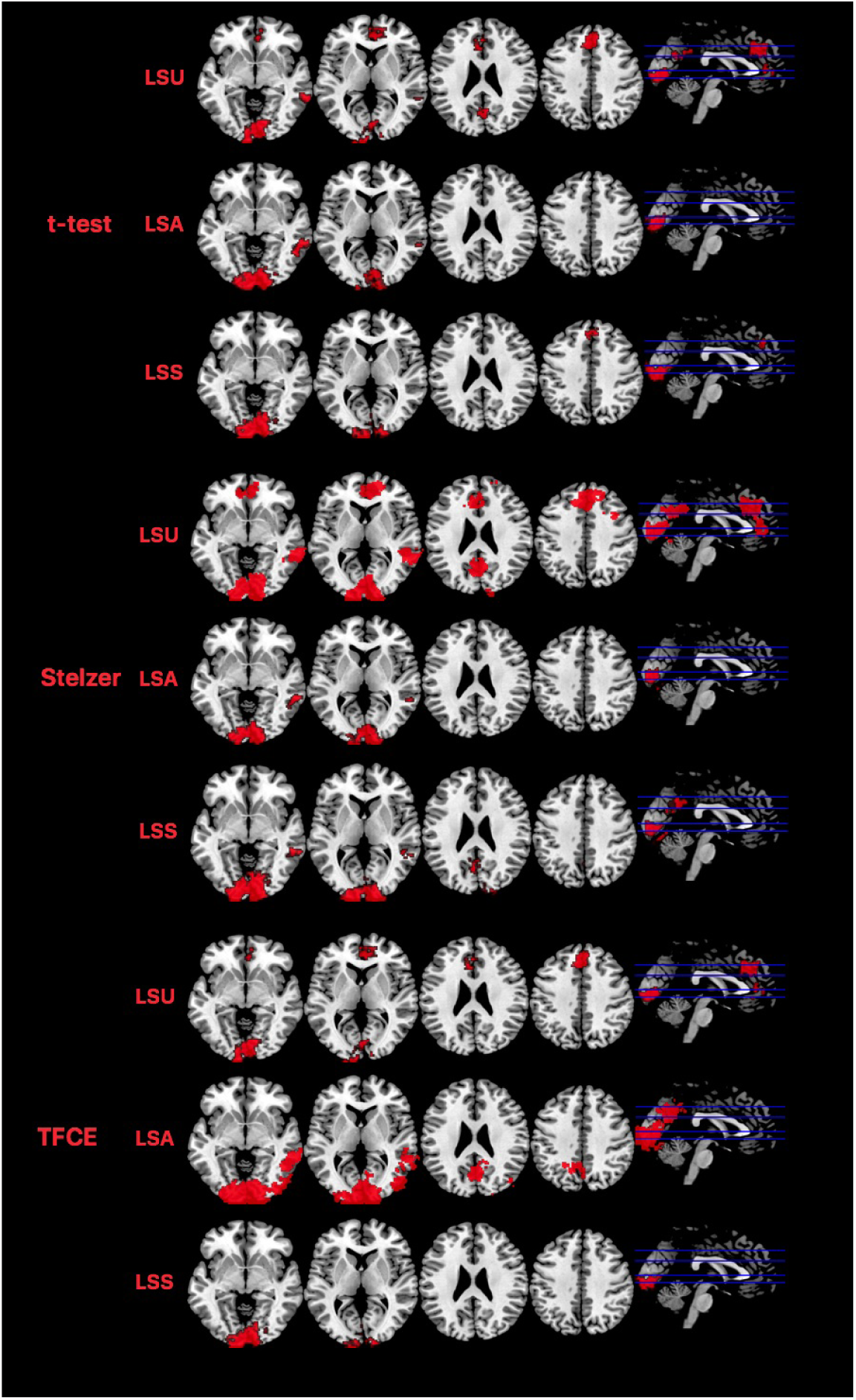
Significant results obtained by the different pattern estimation methods and techniques for evaluating the statistical significance in Dataset 3.

**Figure 17:**
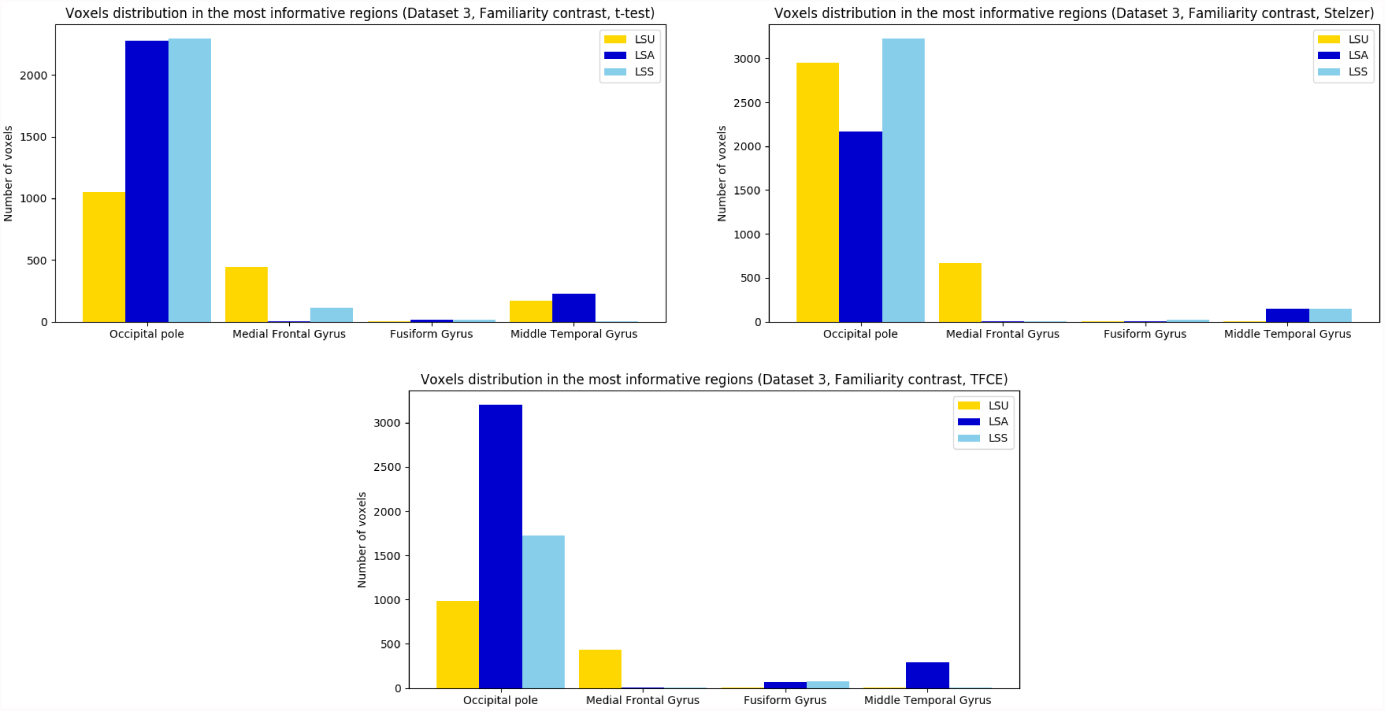
Voxel distribution in the most informative regions for each statistical and pattern estimation method in Dataset 3.

## 4. Discussion

We have shown for the first time that LSS is the most accurate approach for unmixing the contribution to the hemodynamic signal of different events within a trial, regardless of whether they have different or the same duration. Moreover, the non-parametric procedure proposed in Stelzer et al. (2013) is the most sensitive technique when group statistics must be generated from local MVPA approaches such as a searchlight. In this section we will discuss the results obtained by each method (pattern estimation and statistical) for the different datasets evaluated.

### 4.1. Comparison between LSU, LSA and LSS

In Dataset 1, we found large differences in performance across the pattern estimation methods, particularly for the *valence* classification. These differences were present in the two ways of modelling the first event. Estimating responses through LSS allowed us to detect the involvement of a coherent set of brain regions, whereas using LSU and LSA did not yield significant results. Previous studies showed that the performance of LSA and LSS (Abdulrahman and Henson, 2016) is affected by parameters such as the ISI, noise and trial variability. Collinearity is another element that plays a crucial role in the estimation of neural activity. The difficulty of applying decoding analyses in our paradigm is not due to a short interval between consecutive trials, but to the lack of separation between the activity associated with each event within a trial. When the words were modelled as extended events, in accordance to the sustained preparatory activity that they generate, the LSS approach reached its maximal sensitivity. Collinearity cannot be reduced to the way in which events are modelled, as it is also highly affected by the cognitive nature of the process underlying the events. It is worth highlighting that, to the best of our knowledge, this is the first time that these estimation methods are compared in a setting like this. Our results are coherent with findings of previous studies. Analyses carried out by Mumford et al. (2012) concluded that LSS outperforms LSA in high collinearity settings, as it does not employ any regularization strategy. Besides, it is worth remembering that this method was developed due to the poor performance of LSA in rapid event-related designs.

The analyses of the second event of Dataset 1 (e.g. the *fairness* classification) yielded significant results for the three pattern estimation methods, unlike the *valence* classification where only LSS was sensitive enough. Besides, the influence of the different ways of modelling the first event into the estimability of the second event was minimal since results are essentially the same in both contexts. The key of this finding is the classification problem itself. Neural activity differentially associated with valence is hard to obtain, as shown by recent metaanalytic approaches (Lindquist et al., 2015), whereas the fairness of an offer generates large differences and thus it is easier for the LSU approach to make an accurate estimation. Regarding LSA, we mentioned above the large collinearity between the first event (adjective) and the second (offer), so it was highly expected that LSA did not find any informative regions in neither the *valence* nor the *fairness* classification. This raises the intriguing possibility that in contexts where most of the strategies fail to detect differential activity, LSS might be sensitive to small variations.

In Dataset 2 we found large similarities in the results obtained by all pattern estimation methods. A block for each object category was presented only once in each run, which means that no average was applied across experimental conditions of the same type. This yields the same number of beta maps for all classifiers, so that the disadvantages of LSA from a machine learning standpoint are not met. Besides, block settings are not propitious for a better performance of LSS since the overlap of signals is much lower than in event-related designs. Another reason for this similarity is the large perceptual difference in the neural activity elicited by each type of stimulus (faces and houses), so that it is straightforward for a classifier to build a decision hyperplane that properly separates the corresponding activation patterns.

We used Dataset 3 to evaluate the performance of the different pattern estimation methods in a context more similar to our experiment than Dataset 2. In Dataset 3, all pattern estimation methods were able to extract significant regions. Besides, these regions are quite similar regardless of the method used. It is remarkable that LSA allows a good estimation in this setting. There is an important difference in the experimental design that can explain this result: in Dataset 3 all events represent faces: participants evaluate if these faces are familiar/unfamiliar and respond according to that, which involves a brief activity. However, in Dataset 1 participants read an adjective with a certain valence, and according to this valence, they prepare to respond to an offer. Thus, there is a preparatory process that leads to a sustained activity along time.

### 4.2. Comparison between t-test, Stelzer’s and TFCE

As a further goal, we aimed at testing the adequacy of different statistical approaches. For the *valence* classification of Dataset 1, we only obtained significant results when the LSS method was employed, for the two different durations assigned to the first event. The significance maps are essentially the same after applying *t*-test and TFCE, both in the number of significant voxels and in their location. On the other hand, Stelzer’s resulted in a larger sensitivity than the other methods, yielding eight times more significant voxels. Figure 8 compares the uncorrected results for the *t*-test (voxel-level threshold: *p <* 0.001, but uncorrected for multiple comparisons) with the corrected results obtained by Stelzer’s. In this case, there is much more coherence between both methods regarding the number of voxels and, crucially, their location. In fact, the three clusters that Stelzer’s marked as significant are found with the uncorrected *t*-test as well. Therefore, rather than being less sensitive to false positives, Stelzer’s method seems to efficiently detect true data that otherwise do not surpass the statistical threshold. There are several studies that support that non-parametric approaches are able to simultaneously improve the sensitivity while precisely controlling for false positives (e.g. Eklund et al., 2016; Nichols and Hayasaka, 2003; Silver et al., 2011; Stelzer et al., 2013; Winkler et al., 2014). In addition and most interestingly, the largest cluster uncovered by LSS in the *valence* classification resides in the Medial Frontal Cortex (see Figure 9) and includes the peak of maximum differences between positive and negative valence observed in the metaanalyses published by Lindquist et al. (2015) (MNI = [9, 39, −9], see Figure 1). Thus, this close correspondence speaks strongly in favor of the higher sensitivity of the method.

On the other hand, our study is the first to compare Stelzer’s and TFCE methods. Although both use permutation testing for evaluating significance, the way in which they implement permutations may lead to the large differences observed. One of the most appealing aspects of Stelzer’s is that it takes into account the spatial inhomogeneities of the image. In fact, the scheme used by this approach is equivalent to computing a significance threshold for each voxel separately. This controls the false-positives rate in non-informative voxels and avoids being too conservative in the informative ones (Stelzer et al., 2013), which may lend it more sensitive in event-by-event estimations. An encouraging finding is that there is large spatial overlap between the regions that TFCE and Stelzer’s mark as significant. Specifically, all significant voxels in TFCE are also considered significant by Stelzer’s, but the latter adds voxels to the previously identified clusters (see Figure 7). We found even more similarities between Stelzer’s and TFCE in the *fairness* classification. In fact, the way these voxels are distributed is almost identical as Figure 11 reveals. Most information is encoded in the Pre/Postcentral gyrus, the SMA (Supplementary Motor Area) and the Cingulate Gyrus, as Figure 12 shows. These areas are consistent with previous experiments based on the Ultimatum Game (UG), Corradi-Dell’Acqua et al. (2013). For a more detailed explanation of this task and the concordance between the informative regions and our results, see the meta-analysis by Gabay et al. (2014).

As predicted, similarities between the different statistical methods were larger in Dataset 2. Regarding the *t*-test and TFCE, the spatial distribution of the voxels was essentially the same, with a slight boost of 5% in the number of significant voxels when the latter was applied. On the other hand, Stelzer’s yielded 35% more significant voxels, but all the additional ones marked as significant were adjacent to the clusters obtained by the other two methods. Figure 15 highlights the regions where the information is mainly distributed and its variability over different statistical methods, much smaller than in Dataset 1. Results are essentially the same for each pattern estimation and statistical method, principally in the occipital pole and the fusiform gyrus. Stelzer’s yielded more informative voxels in the cerebelum, but the *t*-test and TFCE were more sensitive in the precuneus. It is important to point out the much larger increase in sensitivity that Stelzer’s yielded in Dataset 1 in comparison with Dataset 2. One possibility is that noise was differently distributed in both designs and generated a differential tendency to false-positives. The jitter between experimental conditions in Dataset 1 and the fact that we were isolating different events within a trial may be the reason why a more adequate statistical method leads to larger improvement of sensitivity in this dataset compared to a block design (Dataset 2). We highlight the importance of this finding since although Stelzer’s showed a larger sensitivity in all contexts, it was even higher than the other two methods in the most difficult case, when the overlap and collinearity between conditions were highest. The nature of the classification *per se* may also be of importance in this difference. Whereas the classic block design from Haxby et al. (2001) contrasted two stimuli with large perceptual and phylogenetic differences (e.g. Kanwisher and Yovel 2006), the classification employed in Dataset 1 compared the same physical stimuli (words), equated in length (number of letters), frequency of use and arousal levels. In addition, whereas the brain networks involved in face processing are different from those activated by houses (Haxby et al., 2014), isolating regions with a differential involvement in valence processing is much harder (e.g. Lindquist et al., 2012, 2015).

Results in Dataset 3 show a great similarity between the two methods based on permutations, more sensitive than the *t*-test as in previous datasets. In fact, they are more similar to those obtained in a block-design (Dataset 2) than in the event-related of Dataset 1. Specifically, the occipital pole, followed by the MFG (Medial Frontal Gyrus) and the MTG (Middle Temporal Gyrus) are the most informative regions (see Table 4), which are consistent with the original study (Visconti di Oleggio Castello et al., 2017). It is important to mention that the additional mechanisms that we have employed to ascertain that results in all the analyses conducted are trustworthy. The first one is the proper selection of a searchlight size. Experiments carried out in Etzel et al. (2013) showed that the number of voxels considered informative in a searchlight map tends to grow as the searchlight radius increases, even when the size of the informative region stays fixed. Thus, the larger the searchlight size, the more likely to obtain false positives. This is consistent with findings in Stelzer et al. (2013), where false positives were boosted for a searchlight diameter of 11 voxels. For our analyses, we chose an intermediate value of 8-voxels searchlights to strike a balance between sensitivity and specificity (Arco et al., 2016; Chen et al., 2011). Additionally, we selected a conservative value for the initial-cluster forming threshold to control false positives. The use of a liberal value can have detrimental effects on false positives, location and even interpretation of neural mechanisms (Woo and Wager, 2014). Likewise, Stelzer et al. (2013) fully studied the relationship between this parameter and the results obtained and they highly recommend the election of a *p*-value ranging from 0.005 to 0.001. We chose the most conservative value (*p=0.001*), prioritizing the control of false positives over sensitivity.

## 5. Conclusion

In this work, we compared three different pattern estimation methods, as well as parametric and non-parametric approaches for testing significance in a setting that requires the isolation of a sustained activity from zero-duration events within the same trial. The method with the best performance, Least-Squares Separate (LSS), comprises an iterative fitting of a new GLM for each unique event, which addresses the large overlap of signal from close events. This method was also tested in a block-design and in an event-related design. In both scenarios, this approach demonstrated its ability for improving the sensitivity and provided more information about the brain regions involved in the cognitive process under study. The different results regarding the statistical approach used suggest that using permutation testing in addition to a local-conservative significance threshold indicates that the better performance is due to a better estimation of brain activity and not to an unspecific boost in false-positives. This supports recent claims that the *t*-test is not the proper option to determine the probability of a decoding result at the group level, due to the assumptions about the Gaussianity of the data that are not always met. Our study provides evidence of which method yields a better performance in settings with large collinearity between signals of different duration, which paves the way for future neuroscience studies.

## 6. Acknowledgments

This research is part of J.E.A’s activities for the PhD Program in Information and Communication Technologies of the University of Granada

## References

Abdulrahman, H., Henson, R. N., 2016. Effect of trial-to-trial variability on optimal event-related fMRI design: Implications for Beta-series correlation and multi-voxel pattern analysis. NeuroImage 125, 756–766.

Allefeld, C., Haynes, J.-D., 2014. Searchlight-based multi-voxel pattern analysis of fMRI by cross-validated MANOVA. NeuroImage 89, 345–357.

Arco, J. E., González-Garcfa, C., Ramírez, J., Ruz, M., 2016. Comparison of different methods for brain decoding from fMRI beta maps. Poster presented at 22nd Annual Meeting of the Organization for Human Brain Mapping, Geneve, (Switzerland).

Bode, S., Haynes, J.-D., 2009. Decoding sequential stages of task preparation in the human brain. NeuroImage 45 (2), 606–613.

Brammer, M., Bullmore, E., Simmons, A., Williams, S., Grasby, P., Howard, R., Woodruff, P., Rabe-Hesketh, S., 1997. Generic brain activation mapping in functional magnetic resonance imaging: A nonparametric approach. Magnetic Resonance Imaging 15 (7), 763–770.

Brett, M., Penny, W., Kiebel, S., 2003. An introduction to random field theory. Http://www.fil.ion.ucl.ac.uk/spm/doc/books/hbf2/pdfs/Ch14.pdf.

Bullmore, E. T., Suckling, J., Overmeyer, S., Rabe-Hesketh, S., Taylor, E., Brammer, M. J., 1999. Global, voxel, and cluster tests, by theory and permutation, for a difference between two groups of structural MR images of the brain. IEEE Transactions on Medical Imaging 18 (1), 32–42.

Chen, Y., Namburi, P., Elliott, L., Heinzle, J., Soon, C., Chee, M., Haynes, J., 2011. Cortical surface-based searchlight decoding. NeuroImage 56, 582–592.

Corradi-Dell’Acqua, C., Civai, C., Rumiati, R. I., Fink, G. R., 2013. Disentangling self- and fairness-related neural mechanisms involved in the ultimatum game: an fMRI study. Social Cognitive and Affective Neuroscience 8 (4), 424–431.

Coutanche, M., Thompson-Schill, S., 2012. The advantage of brief fMRI acquisition runs for multi-voxel pattern detection across runs. NeuroImage 61 (4), 1113–9.

Eklund, A., Nichols, T., Knutsson, H., 2016. Cluster failure: Why fMRI inferences for spatial extent have inflated false-positives rate. Proc Nati Acad Sci U S A 113 (28), 7900–5.

Etzel, J., Zacks, J., Braver, T., 2013. Searchlight analysis: promise, pitfalls, and potential. NeuroImage 78, 261–269.

Forman, S., Cohen, J., Fitzgerald, M., Eddy, W., Mintun, M., Noll, D., 1995. Improved assessment of significant activation in functional magnetic resonance imaging (fMRI): use of a cluster-size threshold. Magn. Reson. Med. 33, 636–647.

Friston, K., Fletcher, P., Josephs, O., Holmes, A., Rugg, M., Turner, R., 1998. Event-related fMRI: Characterizing differential responses. NeuroImage 7 (1), 30–40.

Friston, K., Worsley, K., Frackowiak, R., Mazziotta, J., A.C., E., 1994. Assessing the significance of focal activations using their spatial extent. Human Brain Mapping 1, 210–220.

Gabay, A. S., Radua, J., Kempton, M. J., Mehta, M. A., 2014. The ultimatum game and the brain: A meta-analysis of neuroimaging studies. Neuroscience & Biobehavioral Reviews 47, 549–558.

Gaertig, C., Moser, A., Alguacil, S., Ruz, M., 2012. Social information and economic decision-making in the ultimatum game. Front Neurosci 6 (103).

Golland, P., Fischl, B., 2003. Permutation tests for classification: towards statistical significance in image-based studies. Inf. Process. Med. Imaging 18, 330–341.

González-Garcfa, C., Arco, J. E., Palenciano, A. F., Ramfrez, J., Ruz, M., 2017. Encoding, preparation and implementation of novel complex verbal instructions. NeuroImage 148, 264–273.

González-Garcfa, C., Mas-Herrero, E., de Diego-Balaguer, R., Ruz, M., 2016. Task-specific preparatory neural activations in low-interference contexts. Brain Structure & Function 8, 3997–4006.

Hanson, S., Matsuka, T., V Haxby, J., 2004. Combinatorial codes in ventral temporal lobe for object recognition: Haxby (2001) revisited: Is there a “face” area? 23, 156–66.

Haxby, J., Gobbini, M., Furey, M., Ishai, A., Schouten, J., Pietrini, P., 2001. Distributed and overlapping representations of faces and objects in ventral temporal cortex. Science 5539 (293), 2425–30.

Haxby, J. V., Connolly, A., Guntupalli, J. S., 2014. Decoding neural representational spaces using multivariate pattern analysis. Annual review of neuroscience 37, 435–456.

Haynes, J., Rees, G., 2006. Decoding mental states from brain activity in humans. Rev. Neurosci. 7, 523–534.

Hebart, M., Görgen, K., Haynes, J., 2015. The decoding toolbox (TDT): a versatile software package for multivariate analyses of functional imaging data. Frontiers in Neuroinformatics 8, Article 88.

Hebart, M. N., Baker, C. I., 2017. Deconstructing multivariate decoding for the study of brain function. NeuroImage.

Hebart, M. N., Schriever, Y., Donner, T. H., Haynes, J.-D., 2016. The relationship between perceptual decision variables and confidence in the human brain. Cerebral Cortex 26 (1), 118–130.

Henson, R., 2005. Design efficiency in fMRI. URL http://imaging.mrc-cbu.cam.ac.uk/imaging/DesignEfficiency#VII._Should_I_treat_my_trials_as_events_or_epochs_.3F

Kanwisher, N., Yovel, G., 2006. The fusiform face area: a cortical region specialized for the perception of faces. Philos Trans R Soc Lond B Biol Sci 361 (1476), 2109–2128.

Kriegeskorte, N., Goebel, R., Bandettini, P., 2006. Information-based functional brain mapping. Proceedings of the National Academy of Sciences of the United States of America 103 (10), 3863–3868.

Ku, S.-P., Gretton, A., Macke, J., Logothetis, N. K., 2008. Comparison of pattern recognition methods in classifying high-resolution bold signals obtained at high magnetic field in monkeys. Magnetic Resonance Imaging 26 (7), 1007–1014.

Lee, Y.-S., Janata, P., Frost, C., Hanke, M., Granger, R., 2011. Investigation of melodic contour processing in the brain using multivariate pattern-based fMRI. NeuroImage 57 (1), 293–300.

Lindquist, K., Satpute, A., Wager, T., Weber, J., Barrett, L., 2015. The brain basis of positive and negative affect: evidence from a meta-analysis of the human neuroimaging literature. Cereb Cortex 26 (5), 1910–1922.

Lindquist, K., Wager, T., Kober, H., Bliss-Moreau, E., Barrett, L., 2012. The brain basis of emotion: A meta-analytic review. The Behavioral and brain sciences 35, 121–43.

Logothetis, N. K., Wandell, B. A., 2004. Interpreting the bold signal. Annual review of physiology 66, 735–769.

Misaki, M., Kim, Y., Bandettini, P., Kriegeskorte, N., 2010. Comparison of multivariate classifiers and response normalizations for pattern-information fMRI. NeuroImage 53 (1), 103–18.

Moser, A., Gaertig, C., Ruz, M., 2014. Social information and personal interests modulate neural activity during economic decision-making. Frontiers in human neuroscience 8, 31.

Mumford, J. A., Davis, T., Poldrack, R. A., 2014. The impact of study design on pattern estimation for single-trial multivariate pattern analysis. NeuroImage 103 (Supplement C), 130–138.

Mumford, J. A., Turner, B. O., Ashby, F. G., Poldrack, R. A., 2012. Deconvolving bold activation in event-related designs for multivoxel pattern classification analyses. NeuroImage 59 (3), 2636–2643.

Nichols, T., Holmes, A., 2002. Nonparametric permutation tests for functional neuroimaging: a primer with examples. Hum. Brain Mapp. 15 (1), 1–25.

Nichols, T. E., Hayasaka, S., 2003. Controlling the familywise error rate in functional neuroimaging: a comparative review. Statistical Methods in Medical Research 12 (5), 419–446.

Nichols, T. E., Holmes, A. P., 2001. Nonparametric permutation tests for functional neuroimaging: A primer with examples. Human Brain Mapping 15 (1), 1–25.

O’Toole, A. J., Jiang, F., Abdi, H., Penard, N., Dunlop, J., Parent, M., 2007. Theoretical, statistical, and practical perspectives on pattern-based classification approaches to the analysis of functional neuroimaging data. NeuroImage 19 (11), 1735–1752.

Pereira, F., Botvinick, M., 2011. Information mapping with pattern classifiers: a comparative study. NeuroImage 56 (5).

Pereira, F., Mitchell, T., Botvinick, M., 2008. Machine learning classifiers and fMRI: A tutorial overview. NeuroImage 45, S199–209.

Reddy, L., Tsuchiya, N., Serre, T., 2010. Reading the mind’s eye: Decoding category information during mental imagery. NeuroImage 50 (2), 818–825.

Rissman, J., Gazzaley, A., D’Esposito, M., 2004. Measuring functional connectivity during distinct stages of a cognitive task. NeuroImage 23 (2), 752–763.

Russo, F. D., Berchicci, M., Bozzacchi, C., Perri, R., Pitzalis, S., Spinelli, D., 2017. Beyond the bereitschaftspotential: Action preparation behind cognitive functions. Neuroscience & Biobehavioral Reviews 78, 57–81.

Sakai, K., 2008. Task set and prefrontal cortex. Annu. Rev. Neurosci. 31, 219–245.

Silver, M., Montana, G., Nichols, T. E., 2011. False positives in neuroimaging genetics using voxel-based morphometry data. NeuroImage 54 (2), 992–1000.

Smith, S., Nichols, T., 2009. Threshold-free cluster enhancement: Addressing problems of smoothing, threshold dependence and localisation in cluster inference. NeuroImage 44 (1), 83–98.

Sreenivasan, K. K., Curtis, C. E., D’Esposito, M., 2014. Revisiting the role of persistent neural activity during working memory. Trends in Cognitive Sciences 18 (2), 82–89.

Stelzer, J., Chen, Y., Turner, R., 2013. Statistical inference and multiple testing correction in classification-based multi-voxel pattern analysis (MVPA): Random permutations and cluster size control. NeuroImage 65 (Supplement C), 69–82.

Stokes, M., Spaal, E., 2016. The importance of single-trial analyses in cognitive neuroscience. Trends Cogn Sci. 20 (7), 483–6.

Turner, B., 2010. Comparison of methods for the use of pattern classification on rapid event-related fMRI data. In: Annual Meeting of the Society for Neuroscience. San Diego, CA.

Turner, B., Mumford, J., Poldrack, R., Ashby, F., 2012. Spatiotemporal activity estimation for multivoxel pattern analysis with rapid event-related designs. NeuroImage 62(3), 1429–1438.

Visconti di Oleggio Castello, M., Halchenko, Y., Guntupalli, J., Gors, J., Gobbini, M., 2017. The neural representation of personally familiar and unfamiliar faces in the distributed system for face perception. Sci. Rep 7, 12237.

Winkler, A., Ridgway, G., Webster, M., Smith, S., Nichols, T., 2014. Permutation inference for the general linear model. Neuroimage 92, 381–397.

Wolbers, T., Zahorik, P., Giudice, N. A., 2011. Decoding the direction of auditory motion in blind humans. NeuroImage 56 (2), 681–687.

Woo, C.W., K. A., Wager, T., 2014. Cluster-extent based thresholding in fMRI analyses: Pitfalls and recommendations. NeuroImage 91, 412–419.

Worsley, K., Andermann, M., Koulis, T., MacDonald, D., Evans, A., 1999. Detecting changes in nonisotropic images. Human Brain Mapping 8, 98–101.

Worsley, K. J., Evans, A. C., Marrett, S., Neelin, P., 1992. A three-dimensional statistical analysis for CBF activation studies in human brain. Journal of Cerebral Blood Flow & Metabolism 12 (6), 900–918.

